# Multi-day rTMS exerts site-specific effects on functional connectivity but does not influence associative memory performance

**DOI:** 10.1101/2020.04.23.056655

**Authors:** Joshua Hendrikse, James Coxon, Sarah Thompson, Chao Suo, Alex Fornito, Murat Yücel, Nigel C Rogasch

## Abstract

Transcranial magnetic stimulation (TMS) is a non-invasive brain stimulation technique with the capacity to modulate brain network connectivity and cognitive function. Recent studies have demonstrated long-lasting improvements in associative memory and resting-state connectivity following multi-day repetitive TMS (rTMS) to individualised parietal-hippocampal networks. We aimed to assess the reproducibility and network- and cognitive-specificity of these effects following multi-day rTMS. Participants received four days of 20 Hz rTMS to a subject-specific region of left lateral parietal cortex exhibiting peak functional connectivity to the left hippocampus. In a separate week, the same stimulation protocol was applied to a subject-specific region of pre-supplementary motor area (pre-SMA) exhibiting peak functional connectivity to the left putamen. We assessed changes to associative memory before and after each week of stimulation (N = 39), and changes to resting-state functional connectivity before and after stimulation in week one (N = 36). We found no evidence of long-lasting enhancement of associative memory or increased parieto-hippocampal connectivity following multi-day rTMS to the parietal cortex, nor increased pre-SMA-putamen connectivity following multi-day rTMS to pre-SMA. Instead, we observed some evidence of site-specific modulations of functional connectivity lasting ∼24 hours, with reduced connectivity within targeted networks and increased connectivity across distinct non-targeted networks. Our findings suggest a complex interplay between multi-day rTMS and network connectivity. Further work is required to develop reliable rTMS paradigms for driving changes in functional connectivity between cortical and subcortical regions.

## 1. Introduction

Transcranial magnetic stimulation (TMS) is a non-invasive brain stimulation technique with the capacity to modulate brain network connectivity and cognitive function (Fox, Halko, Eldaief, & Pascual-Leone, 2012). A single session of repetitive TMS (rTMS) can induce short-term changes in cortical excitability (Matsunaga et al., 2005) and memory function (Luber et al., 2007) typically lasting less than an hour. Interestingly, recent studies have shown that multiple doses of rTMS delivered over consecutive days may confer longer-lasting (>24 hrs) cumulative effects on brain function and cognitive performance in healthy individuals (Wang et al., 2014; Wang & Voss, 2015). For example, Wang et al. (2014) reported sustained enhancement of associative memory in a sample of 16 healthy individuals (∼24 hours) following five sessions of 20 Hz rTMS applied to subject-specific regions of lateral parietal cortex displaying intrinsic functional connectivity (FC) to left hippocampus. Improvements in memory performance were associated with highly specific increases in FC across the targeted cortico-hippocampal network. These findings suggest that multi-day rTMS may be an effective method of modulating distributed networks critical to learning and memory and commonly implicated in psychiatric (Heckers, 2001; MacQueen et al., 2003) and degenerative disease (Fox et al., 1996). Indeed, such multi-day approaches are closer in design to treatment protocols used in psychiatric disorders such as depression (George & Short, 2014) and therefore may inform future therapeutic applications of stimulation for memory disorders.

The ability to induce long-lasting memory improvement with multi-day stimulation is a prime example of the exciting potential of rTMS. However, the physiological response to non-invasive brain stimulation methods is known to be highly variable between individuals (López-Alonso, Cheeran, Río-Rodríguez, & Fernández-Del-Olmo, 2014), and experimental findings have often proven difficult to replicate (Héroux, Taylor, & Gandevia, 2015). Initial demonstrations of large effects with moderate sample sizes are also prone to effect size overestimation (i.e. ‘The Winner’s Curse’, see Button et al., 2013). Encouragingly, the declarative memory and FC effects initially reported by Wang et al. (2014) following multi-day 20 Hz to the parietal-hippocampal network have since been replicated in similar moderate sample sizes (N = 15 - 16) (Freedberg et al., 2019b; Hermiller et al., 2019). In light of these findings, investigating the reproducibility (i.e. the robustness and generalisability; Milkowski et al., 2018) of these effects with similar multi-day rTMS protocols and a larger sample remains a crucial step in assessing therapeutic potential.

The capacity of rTMS to modulate the activity of targeted brain networks is central to cognitive and clinical applications of stimulation (Shafi, Westover, Fox, & Pascual-Leone, 2012). Whilst the direct effects of TMS are confined to a small area of cortex adjacent to the TMS coil position, indirect effects on FC may be induced across distributed regions on the basis of shared intrinsic FC (Eldaief et al., 2011; Fox et al., 2012). However, recent evidence suggests that the effects of rTMS may vary substantially across different brain regions (Cocchi et al., 2016; Castrillon et al., 2020). Castrillon and colleagues (2020) demonstrated opposing effects of 1 Hz stimulation between sensory and cognitive networks. Stimulation of a frontal cognitive network resulted in widespread decreases in FC, while stimulation of an occipital sensory network produced a paradoxical increase in FC. Further, applying continuous theta burst stimulation to frontal and occipital brain regions also induced opposing effects on FC (Cocchi et al., 2016). Considering that rTMS is applied to different brain networks to treat different psychopathologies (i.e. dorsolateral prefrontal cortex in depression versus orbitofrontal cortex in obsessive-compulsive disorder), investigating the site-specificity of rTMS-induced effects may be an important step in improving treatment efficacy.

The aims of this study were twofold. The first aim was to investigate the reproducibility of long-lasting improvements in associative memory and resting-state connectivity following multi-day 20 Hz rTMS to individualised parietal-hippocampal networks. The second aim was to investigate the site-specificity of changes to FC and associative memory following multi-day rTMS by also targeting a spatially distinct network. We utilised a multi-day 20 Hz rTMS protocol similar to Wang et al. (2014), adopting the same resting-state fMRI target localisation approach and outcome measures (i.e. change in functional connectivity and the face-cued word recall task). Further, we extended the approach of Wang et al. (2014) by using a comparison site, instead of sham rTMS. To do this we applied multi-day 20 Hz rTMS to the pre-supplementary motor area (pre-SMA) to modulate a pre-SMA-putamen network. We chose pre-SMA as a comparison site as it corresponds to a distinct brain network supporting other facets of memory function, namely procedural memory (Doyon et al., 2009).

We expected improvements in associative memory performance, as indexed by the percentage of correctly recalled items on face-cued word recall task, following multi-day parietal stimulation. Further, following multi-day parietal stimulation, we expected an increase in FC across the targeted cortico-hippocampal network consistent with the findings of Wang et al. (2014). We also hypothesised a significant positive correlation between associative memory and FC changes following parietal stimulation. In contrast, following multi-day pre-SMA-stimulation, we expected an increase in FC across the targeted pre-SMA-putamen network with no corresponding improvement in associative memory.

## 2. Methods

### 2.1 Ethics approval

The study was approved by the Monash University Human Research Ethics Committee and all participants provided informed consent prior to their participation. The study conformed to the standards set by the *Declaration of Helsinki* and subjects were remunerated for their participation.

### 2.2 Participants

The sample size was determined from the results of a previous study demonstrating long-lasting changes to cortico-hippocampal functional connectivity and memory performance following multi-day rTMS to left parietal cortex (Wang et al., 2014). The effect size reported by Wang et al. (2014) for memory improvement (Cohen’s *d* = 0.75; derived from differences in raw scores between stimulation and sham conditions), suggests a minimum sample size of 21 for a desired power of 90% at α = .05. Given the tendency for effect sizes to be inflated in the first instance (Button et al., 2013), and the inclusion of an active control condition, we recruited a sample of 40 right-handed healthy adults (52.5% female), aged 25.48 ± 9.35 years (mean ± SD; range 18 – 55). This sample size affords 90% power to detect a moderate effect (*η*^2^_*p*_ = 0.06). Participants reported no contraindications to magnetic resonance imaging or transcranial magnetic stimulation and reported no history of psychological or neurological disorders.

### 2.3 Experimental design

A within-subject cross-over design was used to investigate the effects of multi-day rTMS on associative memory function. Each participant completed two rTMS conditions over the course of two separate weeks; an experimental condition targeting left lateral parietal cortex and another targeting pre-SMA as a comparison site. The stimulation site order was counterbalanced across participants. For comparisons of resting-state network connectivity, MRI scanning was conducted for the first stimulation condition only. Therefore, changes in network connectivity following stimulation to different sites were compared using a between-subjects design (see Figure 1 for overview of experimental design).

**Figure 1.**
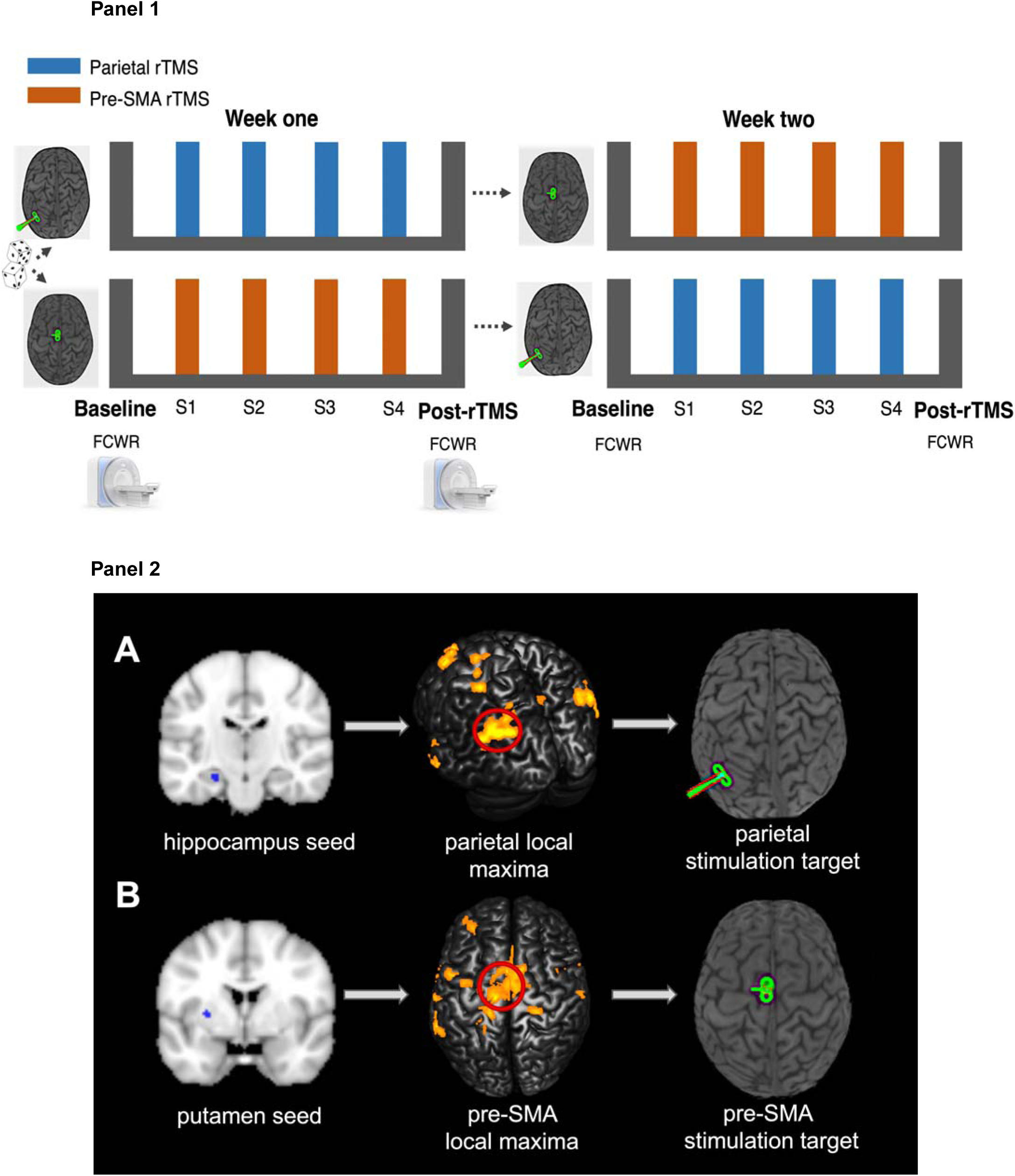
Experimental overview. **The multi-day rTMS protocol is represented in panel 1**. Subjects completed two rTMS conditions across two separate weeks. Each condition included four sessions of personalised rTMS (S1-S4) delivered daily. Stimulation was applied to left lateral parietal cortex functionally integrated within a cortico-hippocampal network in one week, and pre-supplementary motor cortex functionally integrated within a cortico-putamen network in the other week. The order of these conditions was counterbalanced and pseudo-randomised across participants. A minimum one-week break separated the two conditions (range 1-5 weeks). Associative memory performance (face-cued word recall; FCWR) was assessed at baseline (∼1 hour prior to S1) and post-rTMS (∼24 hours following S4). MRI assessments of resting-state functional connectivity were completed in week one at baseline approximately 72 hours prior to S1 and post-rTMS approximately 24 hours after S4. **The rTMS target localisation process is represented in panel 2.** Cortical stimulation sites were determined for each subject on the basis of individual resting-state functional MRI connectivity maps. **(A)** Parietal stimulation sites were derived from a cortico-hippocampal network implicated in associative memory performance. The left hippocampal seed time course was extracted from a 3mm sphere corresponding to the middle of the hippocampus proper, closest to MNI coordinate x = −24, y = −18, z =-18. Parietal targets were defined as the cluster of voxels displaying peak local connectivity within a 15mm radius of MNI coordinate x = −47, y = −68, z = 36 (Wang et al., 2014). **(B)** Pre-SMA stimulation sites were derived from a functionally and spatially distinct cortico-striatal pre-motor network. The left dorso-caudal putamen seed time course was extracted from a 3mm sphere corresponding to MNI coordinate x = −28, y = 1, z = 3. Pre-SMA targets were localised to an area on the cortical surface within left hemisphere, anterior to the anterior commissure (i.e. a positive y MNI coordinate), displaying peak connectivity to the putamen seed. A stereotactic neuronavigation system was used throughout each session to ensure accurate target localisation relative to each subject’s neuroanatomy.

Each condition included daily rTMS sessions delivered over four consecutive days (i.e. Monday – Thursday), with baseline and post rTMS assessments of associative memory performance. Baseline assessments were conducted ∼1.5 hours prior to the first rTMS session (i.e. Monday) and post-rTMS assessments were conducted on Friday, one day after the final rTMS session (parietal condition mean delay = 24.3 ± 2.73 hrs, (mean ± SD; range = 16 −28.5); pre-SMA condition mean delay = 24.2 ± 2.74 hrs, (mean ± SD; range = 18 - 29; *p* = .67). Consistent with the procedure of Wang et al. (2014), each condition was separated by an interval of at least one week (mean = 1.78, range 1 - 5 weeks). The order of experimental versus control weeks was counterbalanced across subjects, and pseudo-randomised to minimise variance associated with gender, age, level of education, self-reported physical activity (IPAQ), and preferred time of day for rTMS administration (i.e. afternoon or morning), using an online algorithm (www.rando.la). Pseudo-randomisation was performed to increase the reliability of between-group comparisons conducted on week one MRI and cognitive data.

The choice to use four as opposed to five days of rTMS (e.g. Wang et al., 2014) was based on scanner availability, with MRI only accessible on weekdays. Instead of conducting our post-rTMS scans on the following Monday, which would have resulted in a 48-72 hour gap following the final rTMS session, we chose to use a shorter four day rTMS protocol and conduct the post scan on Friday with an ∼ 24 hour gap from final rTMS. This stimulation protocol is consistent with other recent studies, which have applied 3 – 4 sessions of rTMS to the parietal-hippocampal network (Freedberg et al., 2019b), and still allowed us to assess the generalisability of stimulation-induced changes to cortical-hippocampal networks and associative memory, although not the direct replicability of the original Wang et al. (2014) protocol.

To investigate functional connectivity following stimulation, resting-state functional MRI scans were conducted on Fridays whenever possible i.e. three days prior to the first rTMS session (interval between baseline MRI and first rTMS session = 75.38 ± 11.03 hrs, (mean ± SD; range = 66.5 - 148) and one day after the final rTMS session (interval between follow-up MRI and final rTMS session = 23.2 ± 3.21 hrs (mean ± SD; range = 18 - 30)).

### 2.4 Associative memory task - Face-cued word recall

To assess the effects of rTMS on associative memory function, we employed the same face-cued word recall task as reported by Wang et al. (2014). Subjects studied a set of 20 human face photographs derived from a database of amateur model headshots (Althoff & Cohen, 1999), presented in greyscale on printed cards. Each card was placed on a table in front of the participant for three seconds, and a unique common English word was read aloud by the experimenter when each card was shown. The words were nouns between 3-8 letters in length, with Kucera-Francis written frequencies between 200 to 2000, and concreteness ratings of 300 to 700 (MRC Psycholinguistic Database; www.psych.rl.ac.uk). Subjects were instructed to memorise the association between the face and word. Following presentation of the 20 face-word pairs, there was a filled delay of approximately one minute involving a procedural task with low cognitive demand (i.e. retrieving an online link to a questionnaire from a personal email inbox). We acknowledge that this filled delay protocol may differ from past studies (Wang et al., 2014; Hermiller et al., 2019). However, this task minimised the potential for participants to employ mnemonic rehearsal strategies during the delay period, which may have led to biased memory assessments. Further, performance on declarative memory tasks (e.g. free word recall) is unaffected by intervening tasks of a semantically distinct nature (Barnes & Underwood, 1959). Hence, we believe that this task with low cognitive demands was unlikely to interfere with associative memory consolidation.’ Following this delay period, the same set of cards was represented to subjects individually in a different randomised order, and subjects were instructed to try and recall the word that accompanied each face. Word recall was scored as correct or incorrect, forgiving errors relating to pronunciation, including plurality.

There were six alternative versions of the task (matched for word concreteness and frequency). Alternative versions were used for baseline and post-rTMS assessments for each stimulation condition (and for a follow-up assessment ∼1.5 weeks following the second condition) and the order of each set was randomised across subjects using a Latin square. To familiarise subjects with the task and minimise practice effects, the remaining sixth version of the task was employed as a practice set, whereby a smaller sub-set of 10 face-word pairs was administered prior to the baseline assessment of the first condition.

### 2.5 Magnetic resonance imaging (MRI)

MRI data were collected from a Siemens 3T Skyra scanner with 32-channel head coil. For both baseline and post rTMS scans, T1-weighted structural images (Magnetization Prepared Rapid Gradient Echo, TR = 2.3 s, TE = 2.07 ms, voxel size 1mm^3^, flip angle 9°, 192 slices) and resting-state Echo Planar Images (TR = 2.46 s, TE = 30 ms, voxel size 3mm^3^, flip angle 90°, 44 slices) with whole brain coverage were acquired. During resting-state scans, subjects were instructed to keep their eyes open and focus on a black fixation cross presented on a white background, whilst trying not to think of anything in particular. MRI scanning was conducted prior to cognitive assessments to ensure that resting-state fMRI measurements were not influenced by completion of cognitive tasks (e.g. to minimise the potential that engaging certain regions during cognitive assessments, such as the hippocampus during associative memory tasks, may alter functional connectivity estimates).

### 2.6 Identification of stimulation locations using resting-state fMRI

Following a similar approach and rationale to Wang et al. (2014), cortical stimulation sites were determined for each subject on the basis of individual resting-state functional MRI connectivity maps calculated from baseline data (Figure. 1). Stimulation sites were chosen on the basis of their superficial location along the cortical surface, allowing direct targeting with TMS, and previously established functional connectivity to distinct cortico-subcortical networks (Di Martino et al., 2008; Wang et al., 2014). Parietal stimulation sites were derived from a cortico-hippocampal network implicated in associative memory performance (Wang et al., 2014). Pre-SMA stimulation sites were derived from a functionally and spatially distinct cortico-striatal pre-motor network (Di Martino et al., 2008) (see Figure. 2)

**Figure 2.**
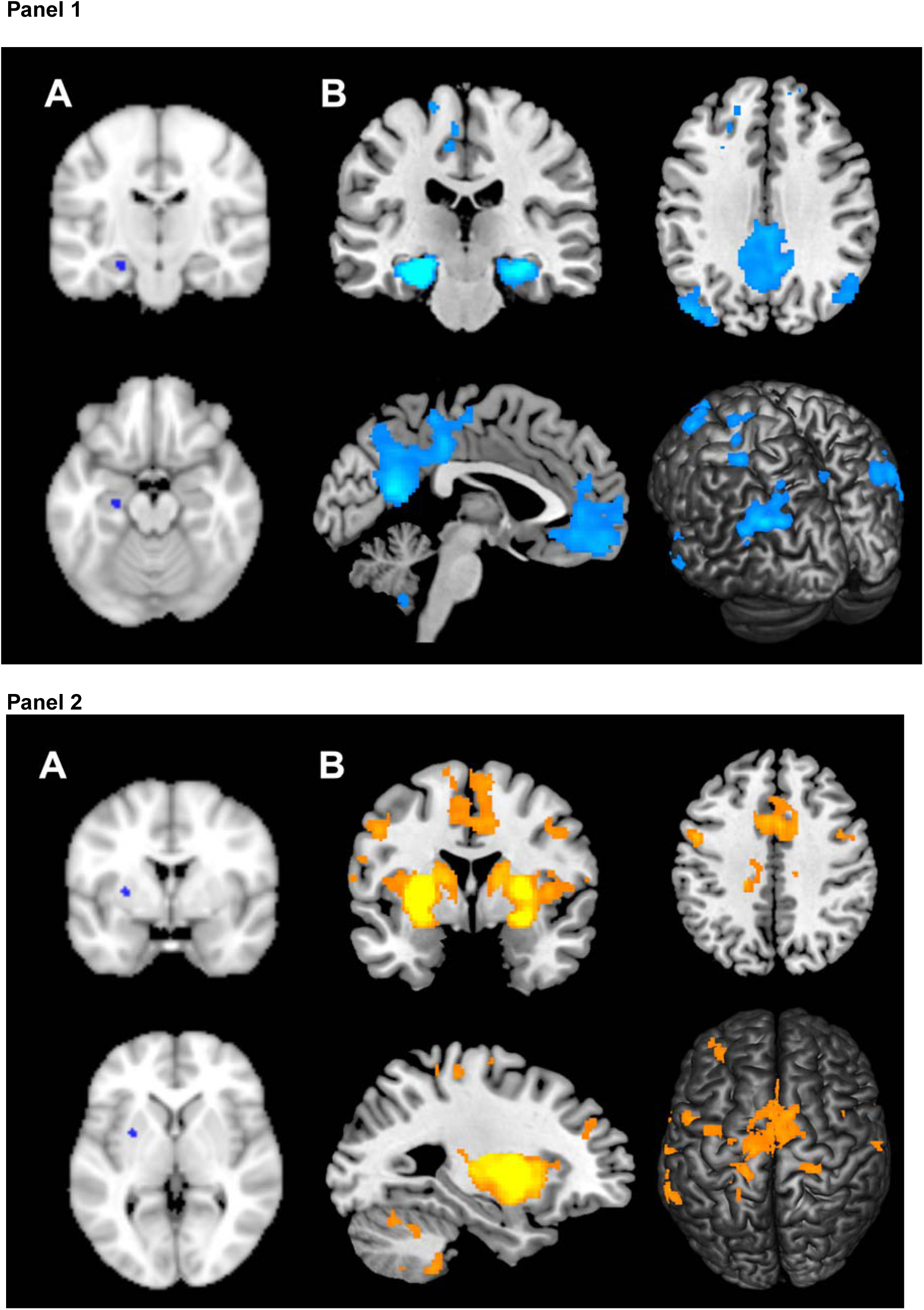
Overview of cortical-hippocampal and cortical-putamen resting-state networks. **Cortical-hippocampal network is represented in panel 1.** To generate functional maps **(B)**, the left hippocampal seed time course was extracted from a 3mm sphere corresponding to middle of hippocampus proper (MNI coordinate x = −24, y = −18, z =-18) **(A)**. This region shows strong functional connectivity to regions within default-mode network and left parietal cortex stimulation target (Wang et al., 2014). Functional maps **(B)** are derived from the group mean of baseline data (voxelwise FWE p < .0001, with extent threshold k > 50 contiguous voxels for illustrative purposes). **Cortical-putamen network is represented in panel 2.** To generate functional maps **(B)**, the left putamen seed time course was extracted from a 3mm sphere corresponding to dorsal putamen (MNI coordinate x = −28, y = 1, z = 3) **(A)**. This region of the putamen shows distinct patterns of connectivity across a pre-motor network, including functional and structural connectivity to the pre-supplementary motor area. Functional maps **(B)** are derived from the group mean of baseline data (voxelwise FWE p < .0001, with extent threshold k > 50 contiguous voxels for illustrative purposes).

A MATLAB-based, in-house pre-processing pipeline was used to determine individualised stimulation targets. Structural and functional images were realigned, normalised to MNI space (MNI_152 template), and co-registered using SPM8. Functional images were slice-time corrected to the first acquired slice and linearly detrended using REST toolbox (Song et al., 2011). Nuisance regressors based on CSF and white matter time courses were generated using CompCor (Behzadi, Restom, Liau, & Liu, 2007), and the fsl_regfilt function (Jenkinson, Beckmann, Behrens, Woolrich, & Smith, 2012) was used to perform motion correction (using six head motion parameters), and nuisance regression of white matter and CSF signals. Functional images were bandpass filtered (0.008 – 0.08 Hz) using REST toolbox (Song et al., 2011) and spatially smoothed using an 8mm Gaussian kernel. An 8 mm smoothing kernel was chosen to increase the signal-to-noise ratio ensuring a reliable functional peak was identified across sites and participants.

To generate baseline functional maps, the time course of two sub-cortical seeds were extracted for each subject using fslmeants function. Specifically, the left hippocampal seed time course was extracted from a 3mm sphere corresponding to the middle of the hippocampus proper, closest to MNI coordinate x = −24, y = −18, z =-18 (Wang et al., 2014). The left dorso-caudal putamen seed time course was extracted from a 3mm sphere corresponding to MNI coordinate x = −28, y = 1, z = 3. This region of the putamen shows distinct patterns of connectivity across a pre-motor network, including functional (Di Martino et al., 2008) and structural connectivity (Alexander, DeLong, & Strick, 1986; Draganski et al., 2008) to the pre-supplementary motor area. To ensure time series reflected fluctuations in grey matter tissue, the extracted seed time courses were weighted by grey matter probability masks derived from each subject’s segmented structural image. The time-series for each seed was used as a regressor to derive individual functional connectivity t maps in SPM8. The group level functional connectivity networks at baseline are shown in Figure 2 (voxelwise FWE p < .0001, with extent threshold k > 50 contiguous voxels for illustrative purposes).

Individual functional connectivity t-maps were used to determine the stimulation targets (Figure 3). Parietal targets were identified from the hippocampus seed-based network and defined as the cluster of voxels displaying peak local connectivity within a 15mm radius of MNI coordinate x = −47, y = −68, z = 36 (Wang et al., 2014) (see supplementary methods for details). Across subjects the average (±SD) MNI coordinate for parietal stimulation was x = - 46.4 (2.9), y = −69.8 (3.7), z = 36.3 (3.8). Consistent with the procedure reported in Wang et al. (2014), stimulation and seed locations were localised to the left hemisphere due to the established role of the left lateral parietal cortex role in associative memory (Wagner, Shannon, Kahn, & Buckner, 2005) and predominant ipsilateral connectivity between the hippocampus and parietal cortex (Kahn, Andrews-Hanna, Vincent, Snyder, & Buckner, 2008). Pre-SMA targets were localised to an area on the cortical surface within left hemisphere, anterior to the anterior commissure (i.e. a positive y MNI coordinate), displaying peak connectivity to the putamen seed. Pre-SMA stimulation was biased towards left hemisphere to ensure consistency between the two stimulation conditions (i.e. ipsilateral relationship between sub-cortical seed and stimulation site). Across subjects the average (±SD) MNI coordinate for pre-SMA stimulation was x = −4.6 (2.7), y = 2.9 (2.4), z = 69.7 (4.1), conferring with past studies that have localised pre-SMA to 1-4cm anterior to the scalp vertex (Hamada, Ugawa, & Tsuji, 2009; Matsunaga et al., 2005) (see supplementary figures S1 - S3 for overlay of individualised seed and stimulation locations, and corresponding resting-state networks).

**Figure 3.**
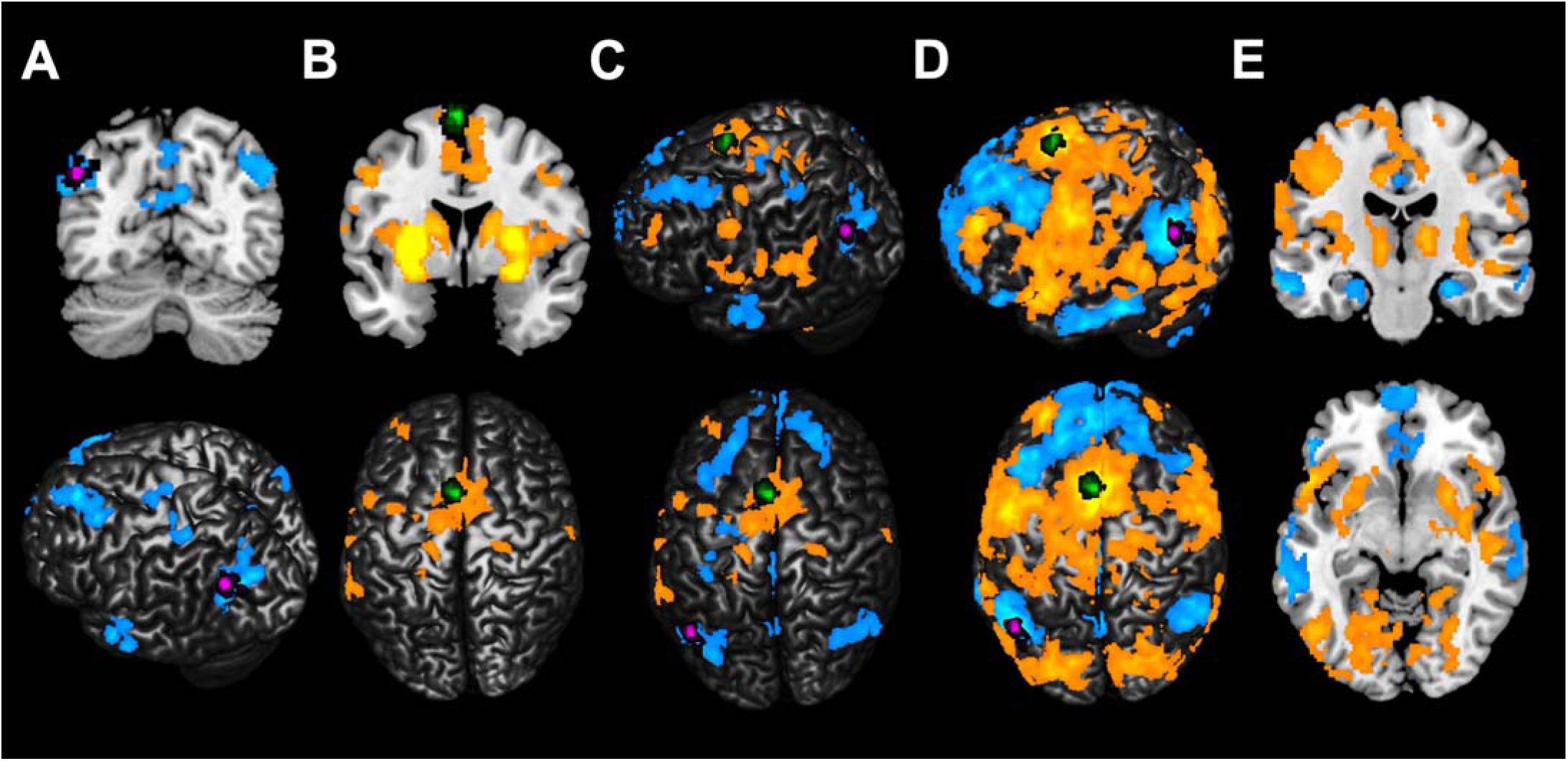
Overlay of individualised stimulation targets on resting-state networks. **(A) Individual parietal stimulation targets & cortico-hippocampal resting-state network.** Parietal stimulation targets (represented in magenta) were selected within a 15mm radius of MNI coordinate x = −47, y = −68, z = 36. Stimulation targets are localised to region of the left parietal cortex functionally connected to left hippocampus. The spatial overlap of stimulation targets across subjects is shown, with greater proportion of overlap represented by lighter magenta colouration. **(B) Individual pre-SMA stimulation targets & cortico-putamen resting-state network.** Pre-SMA targets (represented in green) were localised to an area on the cortical surface within left hemisphere, anterior to the anterior commissure (i.e. a positive y MNI coordinate). Stimulation targets are localised to region of pre-SMA functionally connected to left putamen. The spatial overlap of stimulation targets across subjects is shown, with greater proportion of overlap represented by lighter green colouration. **(C-E) Distinct spatial profile of targeted resting-state networks.** Cortical-hippocampal network is represented in blue-light blue colour scale and shows functional connectivity across default-mode network. Cortico-putamen network is represented in yellow-orange colour scale and shows distinct pre-motor functional connectivity. **C)** Resting-state networks extracted from sub-cortical targets (i.e. hippocampus & putamen). **D-E)** Resting-state networks extracted from cortical stimulation sites (i.e. parietal cortex & pre-SMA), shown across the cortical surface **(D)** and between sub-cortical targets **(E)**. Seeding either the cortical stimulation site or corresponding sub-cortical target extracted a spatially consistent network. Parietal stimulation targets are displayed in magenta and pre-SMA targets are displayed in green (voxelwise FWE p < .0001, with extent threshold k > 50 contiguous voxels for illustrative purposes).

### 2.7 Repetitive transcranial magnetic stimulation

At the beginning of each rTMS session, resting-motor threshold was determined using electromyography recorded from the right first dorsal interosseous (FDI) muscle. Biphasic pulses were applied to the left primary motor cortex (M1) using a MagVenture MagPro X100 stimulator and B-65 figure-8 cooled coil (75mm outer diameter). The motor ‘hotspot’ was determined as the area of scalp that produced the largest and most consistent motor-evoked potential in the targeted FDI muscle, and resting motor threshold was defined as the minimum stimulation intensity necessary to evoke a potential with a peak-to-peak amplitude of ≥ 50 µV in the resting FDI in at least 5 out of 10 consecutive trials.

For both stimulation conditions, rTMS was administered at 100% of resting-motor threshold at 20 Hz (2s on, 28s off) for 20 minutes (1600 pulses total) daily for four consecutive days (see table 1 for maximum stimulator intensity values). These parameters conform to internationally established safety guidelines (Rossi, Hallett, Rossini, & Pascual-Leone, 2009) but as an additional precautionary measure, EMG recordings from the FDI muscle were continuously monitored throughout rTMS for evidence of altered motor activity changes (i.e. kindling). A stereotactic neuronavigation system was used throughout each session, enabling real-time monitoring of the TMS coil position to ensure accurate target localisation relative to each subject’s neuroanatomy (Brainsight, Rogue Research; with the exception of subjects 1-9 for whom a Zebris system was used). Analysis of associative memory change was performed for the sub-sample that completed the protocol with the Brainsight system (N = 31) (see supplementary materials), however the results were consistent with the complete sample (N = 40), reported herein. Stimulation targets were loaded onto each subject’s normalised T1-weighted image and aligned to the middle of the gyral crown to maximise the field size induced in cortical grey matter by the TMS pulse (Thielscher, Opitz, & Windhoff, 2011). For the parietal condition, stimulation was delivered to the target site with the coil handle perpendicular to the long axis of the gyrus to induce posterior/anterior current flow. For the pre-SMA condition, the coil was held with the coil handle pointing left, i.e. perpendicular to the midsagittal plane. During stimulation, subjects were seated in an upright position with neck support and instructed to minimise head movement. Coil position relative to the rTMS target site was continuously monitored by the experimenter holding the coil, and immediately corrected following any inadvertent head movement.

**Table 1:** average TMS intensities. Percentage of maximum stimulator output (MSO) (MagVenture X100, Biphasic) at which rTMS was administered (shown for within-subject comparisons, and week one stimulation groups). Parentheses indicate standard error of the mean.

|  | Within-subjects<br>cross-over<br>(N = 39) | Parietal stimulation<br>Week 1<br>(N = 18) | Pre-SMA stimulation<br>Week 1<br>(N = 18) |
| --- | --- | --- | --- |
| TMS intensity (% MSO) | 47.41% (1.30) | 46.33% (1.95) | 48.44% (1.78) |

### 2.8 Data analysis

#### 2.8.1 Face-cued word recall task analysis

For one subject, scheduling conflicts meant that it was not feasible to conduct post-rTMS assessments 24 hours following the final rTMS session. Thus, data from this subject was removed and analyses of associative memory change were conducted on N = 39. Performance on the face-cued word recall task was scored as the percentage of correctly recalled words out of a total of 20. Individual z scores more extreme than 2.58 were labelled as outliers and removed from subsequent analysis (as specified in the results where applicable). To investigate the effects of parietal stimulation on associative memory performance, a 2 x 2 repeated measures ANOVA was conducted with within-subject factors of stimulation condition (parietal, pre-SMA) and time (baseline, post-rTMS). A paired-sample t-test was also conducted on the follow-up assessment data to investigate whether stimulation elicited long-lasting (∼1.5 week) changes to associative memory performance (Wang & Voss, 2015). In order to account for inter-subject variability in task performance, an additional paired-sample t-test (Wang et al., 2014) was also conducted on Δ values for each stimulation condition expressed as: Δ memory = (post-rTMS % correct – baseline % correct) / baseline % correct.

#### 2.8.2 Resting-state fMRI analysis

Previous lines of evidence have demonstrated that pre-processing methods, in particular denoising methods to minimise motion-related artefact, can exert considerable influence on resting-state fMRI group-level comparisons (Parkes, Fulcher, Yücel, & Fornito, 2018). Therefore, data were pre-processed using two separate pre-processing pipelines, including the previously described MATLAB-based, in-house pipeline, and fMRIprep (version 1.1.1), a ‘best-in-breed’ workflow developed to ensure high-quality outputs, robust to data idiosyncrasies (Esteban et al., 2019). fMRIprep was run with default parameters, including ICA-AROMA denoising and susceptibility-derived distortion estimation (AFNI 3dqwarp). The choice of pre-processing pipeline did not significantly alter second-level group comparisons, thus fMRIprep was employed for the final reported analysis.

For each network of interest, we examined changes to FC from both the site of stimulation and corresponding sub-cortical target. To investigate changes to cortical-hippocampal FC, we extracted the time course of subject-specific seed coordinates corresponding to the parietal stimulation site and corresponding left hippocampus target. To investigate changes to cortical-putamen FC, we extracted the time course of subject-specific seed coordinates corresponding to the pre-SMA stimulation site and corresponding left putamen target. Time courses were extracted from the pre-processed functional image using fslmeants for baseline and post-rTMS sessions. Time series were weighted by individual’s grey matter probability mask and resulting time series were entered into first-level analysis in SPM8 to generate individual contrast maps.

Second-level group comparisons of FC change were conducted using SPM and run separately for each seeded region to eliminate statistical issues relating to multi-collinearity.

T maps were converted to Fisher’s Z using the SPM8 function imcalc. Similar to the analytical approach of Wang et al. (2014), difference maps were calculated by subtracting the baseline MRI assessment from the post-rTMS assessment and compared between stimulation conditions using an independent-samples t-test design. Additionally, to account for the possibility that stimulation may have altered FC across broader networks, within-subject comparisons were also conducted for each stimulation condition. Specifically, change to parietal and hippocampal FC (seed in the parietal stimulation site and hippocampus) were analysed following parietal stimulation, and changes to pre-SMA and putamen FC (seed in the pre-SMA stimulation site and putamen) were analysed following pre-SMA stimulation, using a paired-sample t-test design (baseline vs post-stimulation). Additionally, to examine network-specific changes following stimulation we conducted a spatially constrained analysis between stimulation sites and corresponding sub-cortical targets (see supplementary materials for details).Group-level connectivity change maps were FWE-corrected at the cluster level with a cluster defining height threshold of p < .05 to approximate the approach of Wang et al. (2014). Given subsequent evidence that more stringent primary thresholds are more appropriate (Eklund, Nichols, & Knutsson, 2016), we also report results using a more stringent cluster defining height threshold of p < .001, which is commonly used for cluster level FWE inference in SPM.

#### 2.8.3 Parietal-hippocampal functional connectivity and associative memory

Connectivity change between stimulation target and sub-cortical seed was calculated in SPM8, using Marsbar toolbox (Brett, Anton, & Valabregue, 2002). We extracted beta values between subject-specific hippocampal locations (3mm spherical seed) and the stimulation site within left parietal cortex (10 mm spherical seed, approximating the estimated e-field diameter induced by the TMS pulse (De Geeter, Crevecoeur, Leemans, & Dupré, 2015)), centred on MNI coordinate x = −41, y = −67, z = 36. To examine the relationship between changes in resting-state connectivity and memory performance, Pearson’s correlations (two-tailed, p < .05) were conducted between change scores calculated from parietal-hippocampal connectivity beta weights (Δ = post beta– pre beta) and Δ memory. This analysis included only those subjects who received parietal stimulation in the first condition.

## 3. Results

Data were collected from a sample of 40 participants (19 males, 21 females) with an average age of 25.48 (SD ± 9.35) years. Overall, both conditions of TMS were well tolerated by subjects. Two participants reported transient mild headaches following rTMS (one following parietal stimulation and one following pre-SMA stimulation), but no other adverse events were reported.

TMS intensities (% of maximum stimulator output) for the complete cross-over sample, and week one stimulation groups (used for FC comparisons) are reported in table 1. There was no significant difference in TMS intensity between week one parietal and pre-SMA stimulation groups (t_1,34_ = −0.80, p = .43, Cohen’s d = 0.27).

Due to scheduling conflicts, 2/40 subjects did not complete the MRI protocol. Data were also omitted from an additional two subjects (due to error in MRI console software (n = 1) and excessive head motion (n = 1) during acquisition (frequent framewise displacement > 2 mm, max = 4.73 mm; Parkes, Fulcher, Yücel, & Fornito, 2018). Thus, MRI analyses were conducted with N =36, whereby comparisons were made between individuals who received parietal stimulation (n = 18) versus pre-SMA stimulation (n = 18) in week one (see figure 3). There were no significant differences between groups in regards to age (parietal stimulation = 24.89 ± 9.05 years; pre-SMA stimulation = 24.83 ± 8.90 years, t_1,34_ = 0.02, p = .99, Cohen’s d = 0.00), or sex (parietal stimulation = 50.0 % female; pre-SMA = 61.1 % female, (1, N = 36) = 0.45, p = .50, Cramer’s V = .11).

### 3.1 Face-cued word recall task

Based on the findings of Wang et al. (2014, 2015), we hypothesised significant improvements in face-cued word recall accuracy following parietal stimulation, relative to stimulation of the pre-SMA active control site. Contrary to our expectations, a within-subject 2 x 2 repeated-measures ANOVA revealed no significant interaction between Stimulation Condition and Time (*F*_1,38_ = 0.02, *p* = .89, *η*^2^_*p*_ = .00), no main effect of Time (baseline, post-rTMS; F_1,38_ = 0.35, p = .56, *η*^2^_*p*_ = .01), and no main effect of Stimulation Condition (parietal, pre-SMA; F_1,38_ = 1.86, p = .18, *η*^2^_*p*_ = .05).

Two additional paired-sample t-tests were conducted to assess changes in associative memory performance. Associative memory estimates taken at follow-up (i.e. ∼1.5 weeks following final rTMS session) were compared across stimulation conditions. No significant differences were found between parietal stimulation (M = 45.13, SD = 25.43) and pre-SMA stimulation (M = 43.59, SD =23.87), t_1,38_ = 0.59, p = .56, Cohen’s d = 0.09. A paired-sample t-test was also conducted on Δ values calculated for each stimulation condition. Two outlier values (Z-score > 2.58) were identified from the pre-SMA stimulation condition, and these datapoints were removed from the analysis. No significant differences were found between Δ parietal stimulation (M = 0.03, SD = .43) and Δ pre-SMA stimulation (M = 0.02, SD = 0.46), t_1,36_ = 0.09, p = .93, Cohen’s d = 0.01.

Supplementary analyses were also conducted to address potential confounding variables. To account for possible carryover effects of pre-SMA stimulation on the response to parietal stimulation or associative memory estimates, we compared baseline and post-rTMS associative memory scores in subjects who received parietal stimulation in the first week of the study (N = 20). There were no significant differences in associative memory present in this sub-sample (p = 1.00, Cohen’s d = 0.00; see supplementary materials for details). Additionally, to account for the possibility that young and middle-aged adults may have responded differently to multi-day rTMS, an additional analysis of associative memory was performed on the young adult sample (N = 35, aged 18 – 31 years). There were no significant differences between parietal and pre-SMA stimulation conditions in this sub-sample (p = .65, Cohen’s d = 0.08; see supplementary materials for details). Overall, these results are consistent with those reported from the complete sample.

Given that these results do not align with those previously reported in the literature, we conducted Bayesian paired-sample t-tests on associative memory estimates using JASP (v 0.10.2) to quantify the relative evidence for the alternative vs. the null hypothesis. Based our hypothesised increase in associative memory performance following parietal stimulation, the prior was set in support of the alternative over null hypothesis (i.e. BF_10_). The Cauchy parameter was set to a conservative default value of 0.707 (Ly, Verhagen, & Wagenmakers, 2016; Rouder, Speckman, Sun, Morey, & Iverson, 2009). Percentage correct values before and after parietal stimulation were analysed using a model based on a non-directional alternative hypothesis, and results showed moderate evidence in favour of the null hypothesis (BF_10_ = 0.20). The Δ values calculated for each stimulation condition were analysed using a non-directional model and also showed moderate evidence in favour of the null hypothesis (BF_10_ = 0.18). In summary, our data does not support the hypothesis that multi-day parietal stimulation improves associative memory.

### 3.2 Resting-state fMRI

#### 3.2.1 Parietal stimulation

We first compared changes in connectivity seeded from the hippocampus following parietal stimulation (within-subject comparisons; baseline vs post-rTMS). No significant clusters surviving an FWE-corrected p < .05 cluster threshold were observed. However, when directly comparing changes in hippocampal seeded connectivity between the parietal and SMA stimulation groups (between-subject comparisons; post-pre between conditions) we found evidence for divergent effects between sites, with parietal stimulation tending to decrease connectivity between the hippocampus and paracingulate cortex, left putamen, temporal fusiform cortex and postcentral gyrus, whereas SMA stimulation tended to increase connectivity between these regions (FWE-cluster corrected p < .05) (see figure 5). These effects did not survive FWE-cluster correction using a p < .001 cluster-forming threshold.

**Figure 4.**
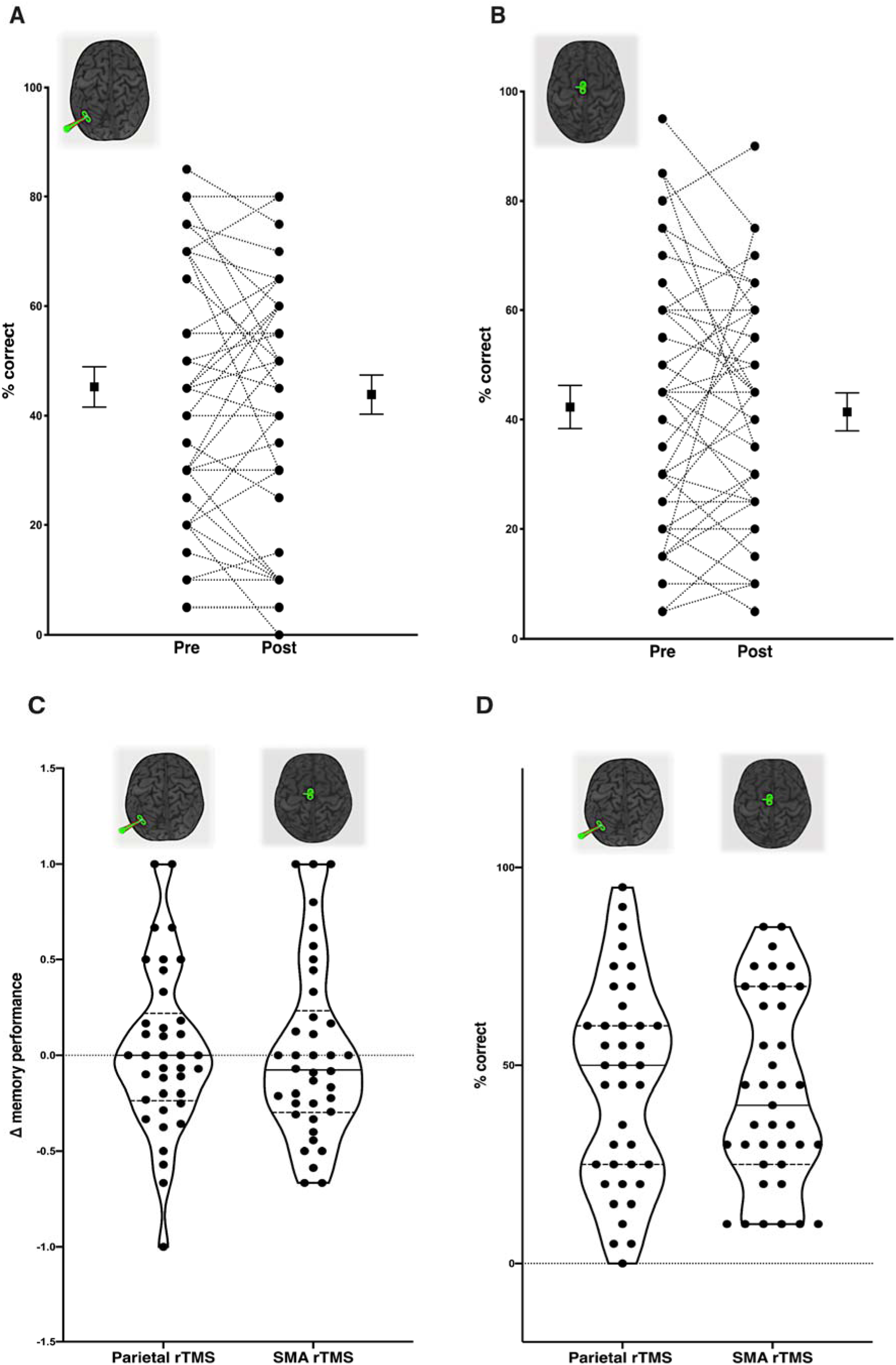
Assessments of associative memory performance. No evidence of sustained improvement in associative memory was shown following parietal or pre-SMA active control stimulation. **A**) Associative memory performance following parietal stimulation (circles represent individual scores; squares and error bars depict group mean and standard error); **B)** Associative memory performance following active control pre-SMA stimulation; **C)** Change in associative memory following parietal stimulation (left) and pre-SMA stimulation (right) expressed as change relative to baseline performance. **D)** Comparison of associative memory performance at follow-up (∼1.5 after final session) following parietal stimulation (left and pre-SMA stimulation (right).

**Figure 5.**
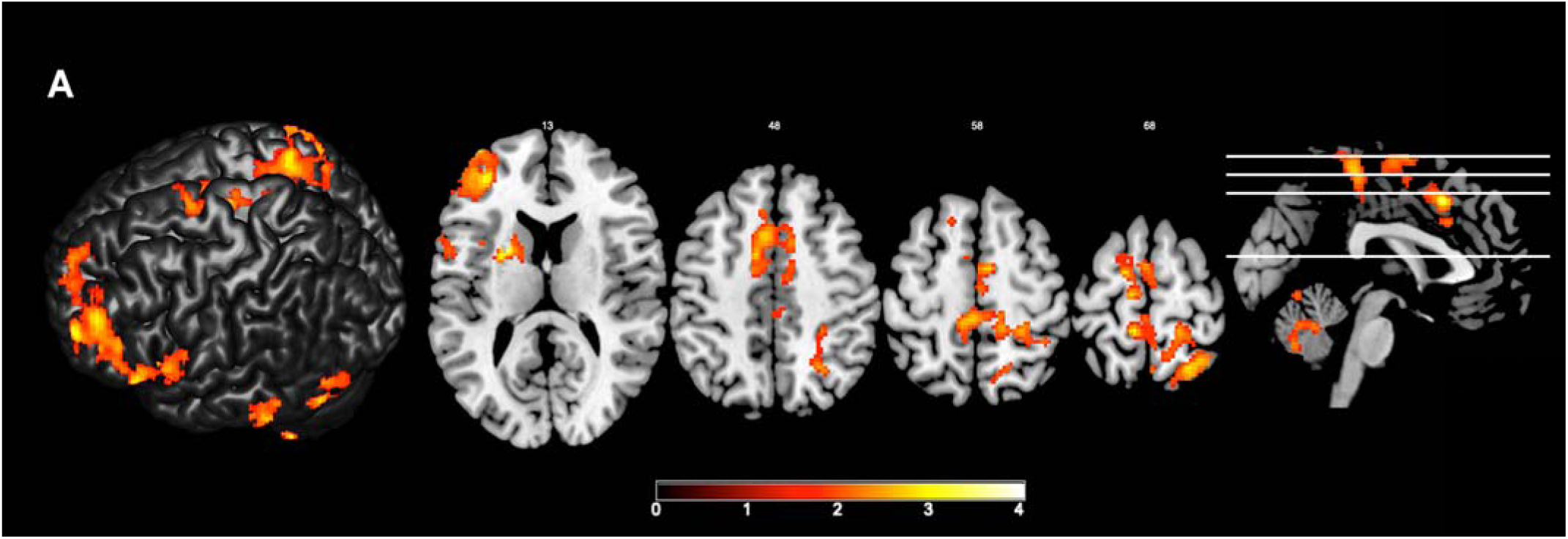
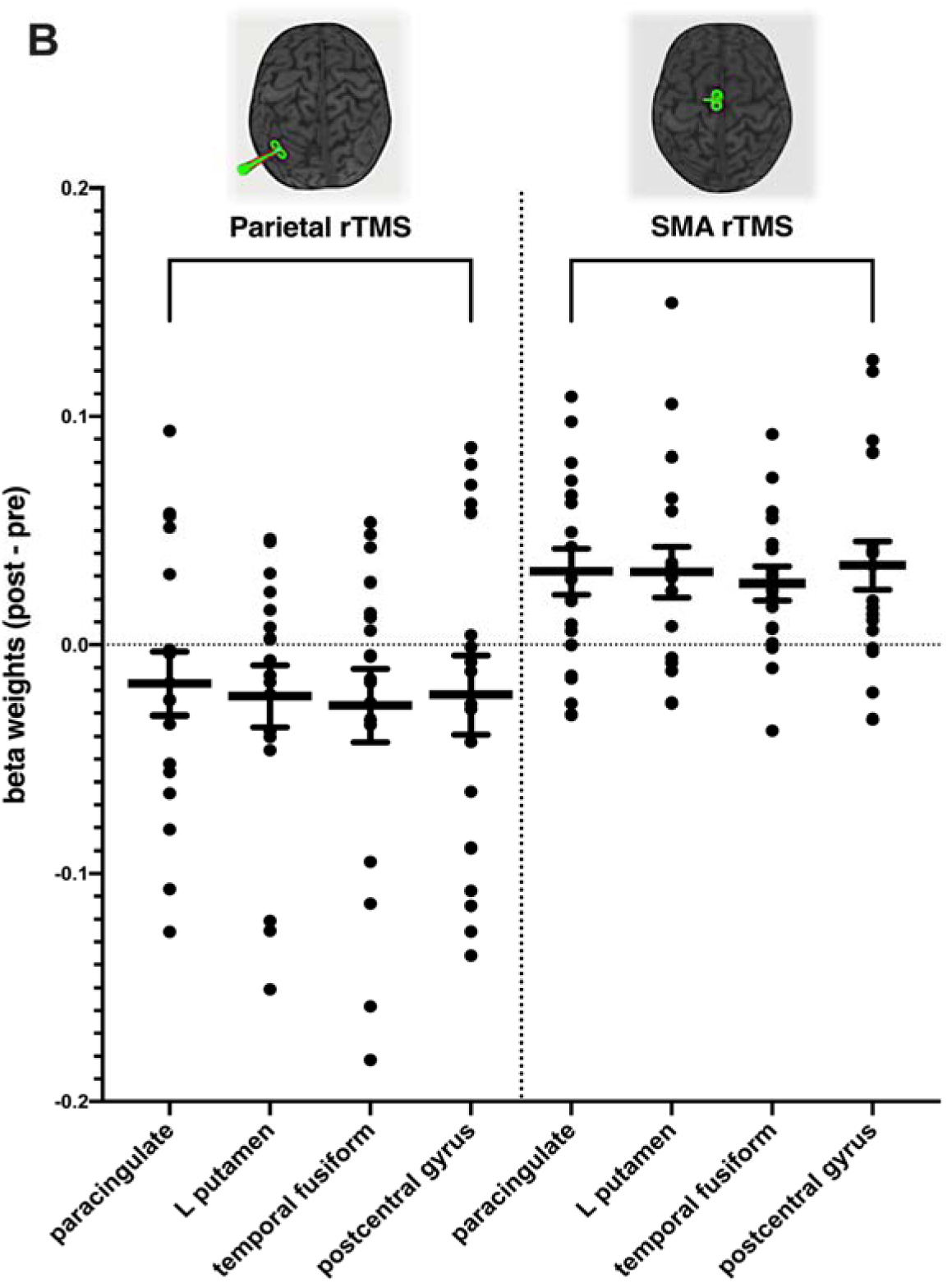
Resting-state connectivity change from hippocampus seed. Evidence of differential effects on hippocampal connectivity between multi-day parietal and pre-SMA stimulation. Subtle evidence of reduced FC following parietal stimulation and increased FC following pre-SMA stimulation between hippocampus seed region and locations outside of *a priori* networks of interest. **A)** Regions showing altered hippocampal FC following stimulation, including paracingulate cortex, left putamen, temporal fusiform cortex and postcentral gyrus, overlaid on anatomical template. **B)** Beta weights obtained from regions showing significant change in FC with the hippocampus seed region following stimulation (between-subject comparison; post-pre between conditions). Left side of plot depicts changes to beta weight strength following parietal stimulation, with evidence of net reduction in functional connectivity. Right side of plot depicts changes to beta weight strength following pre-SMA stimulation, with evidence of net increase in FC. Error bars reflect mean and standard error.

We also repeated our analyses without CSF and WM nuisance regression in line with past studies (Freedberg et al., 2019b, Wang et al., 2014), however there were no significant clusters surviving an FWE-corrected p < .05 cluster threshold for both within-subject (baseline vs post-rTMS, parietal stimulation) and between-subject (post-pre between conditions) comparisons.

As we used individualised cortical targets to determine the stimulation site, we also directly seeded the targeted cortical regions to compare changes in connectivity following stimulation. Contrary to our expectations, parietal stimulation resulted in subtle increases in FC between the stimulation site and occipital cortex (within-subject comparisons; baseline vs post-rTMS). When comparing changes between the parietal and SMA stimulation groups (between-subject comparisons; post-pre between conditions), we again found evidence of divergent effects, with parietal stimulation tending to decrease FC between stimulation site and left medio-temporal cortex close to left hippocampus, whereas SMA stimulation resulted in a net increase in FC between these regions (FWE-cluster corrected p < .05) (see figure 6). These effects did not survive FWE-cluster correction using a p < .001 cluster-forming threshold.

**Figure 6.**
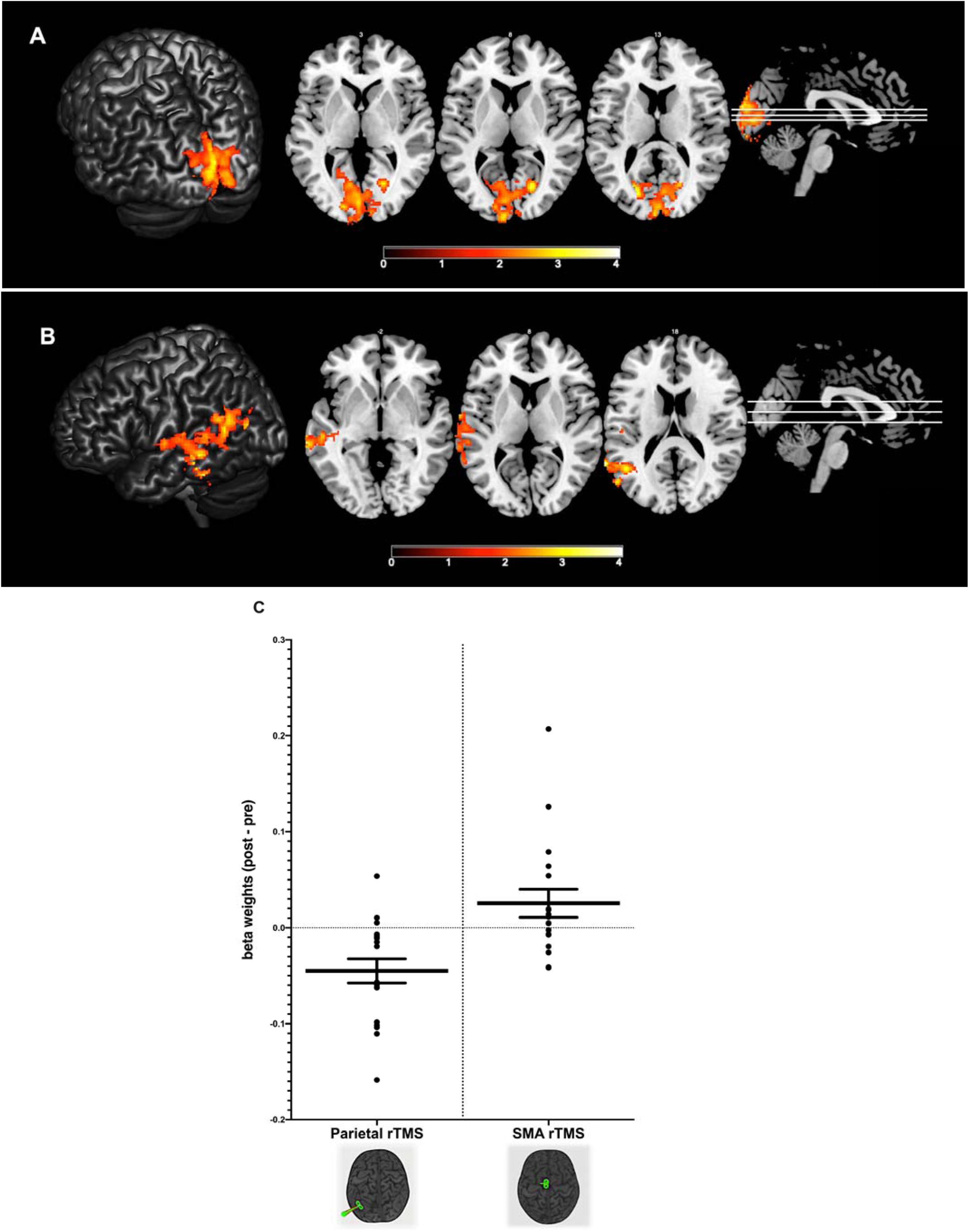
Resting-state connectivity change from parietal stimulation site seed. **A)** Increased FC between parietal seed region and occipital cortex following parietal stimulation. **B)** Evidence of differential effects on parietal connectivity between parietal and pre-SMA stimulation. Subtle evidence of reduced FC following parietal stimulation and increased FC following pre-SMA stimulation in left temporal regions close to hippocampus. **C)** Beta weights obtained from regions showing significant change in FC with the parietal seed region following stimulation (between-subject comparison; post-pre between conditions). Left side of plot depicts changes to beta weight strength following parietal stimulation, with evidence of net reduction in functional connectivity. Right side of plot depicts changes to beta weight strength following pre-SMA stimulation, with evidence of net increase in functional connectivity. Error bars reflect mean and standard error.

Taken together, we could not find any evidence that multi-day parietal stimulation increased connectivity in a parieto-hippocampal network. If anything, our data indicates that parietal stimulation tends to reduce connectivity in hippocampal networks, including connectivity with the stimulation site.

#### 3.2.2 Pre-SMA stimulation

We next compared changes in connectivity seeded from the putamen following pre-SMA stimulation. We found evidence of increased FC between putamen and left frontal cortex (FWE-corrected p < .05; within-subject comparisons; baseline vs post-rTMS; see figure 7), though these effects did not survive FWE-cluster correction using a p < .001 cluster-forming threshold. We found no evidence of FC change when directly comparing changes in putamen seeded connectivity between the parietal and SMA stimulation groups (between-subject comparisons; post-pre between conditions).

**Figure 7.**
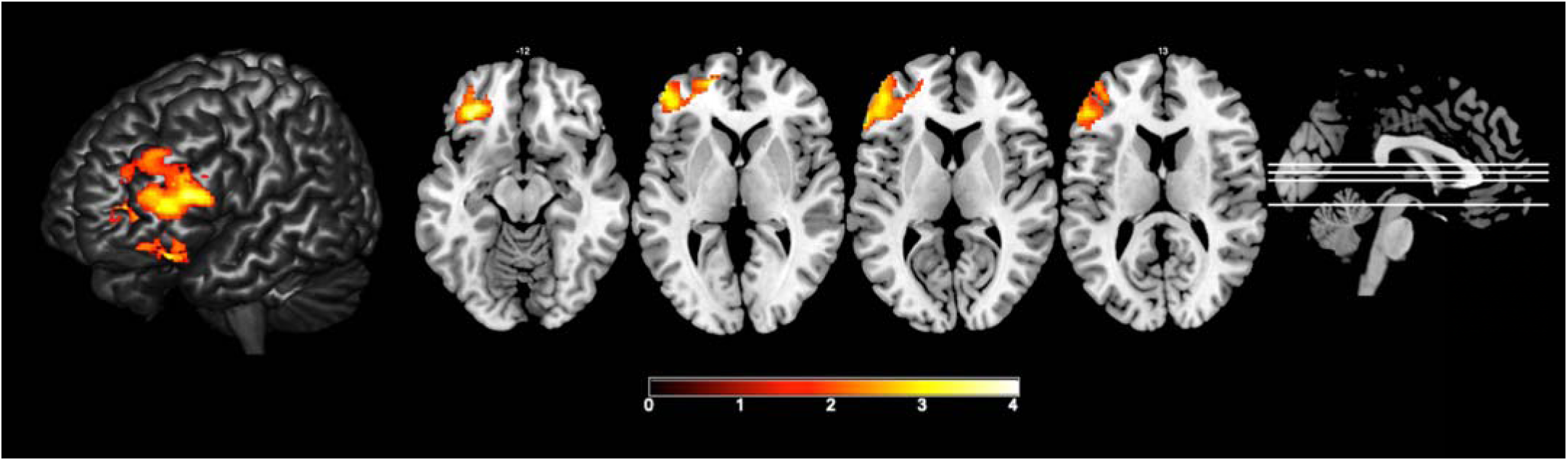
Resting-state connectivity change from putamen seed. Evidence of increased FC between putamen seed region and left frontal cortex following pre-SMA stimulation. No differences in FC were observed when comparing between parietal and pre-SMA stimulation.

We also directly seeded the individualised pre-SMA stimulation target to compare changes in connectivity following stimulation. Contrary to our expectations, we found no evidence of increased FC following pre-SMA stimulation (FWE-cluster corrected p < .05; within-subject comparisons; baseline vs post-rTMS; see figure 8). When comparing changes between the parietal and SMA stimulation groups (between-subject comparisons; post-pre between conditions), we found evidence of decreased FC between stimulation site and occipital cortex following pre-SMA stimulation, and increased FC across these regions following parietal stimulation (FWE-cluster corrected p < .05) (see figure 8). These effects did not survive FWE-cluster correction using a p < .001 cluster-forming threshold.

**Figure 8.**
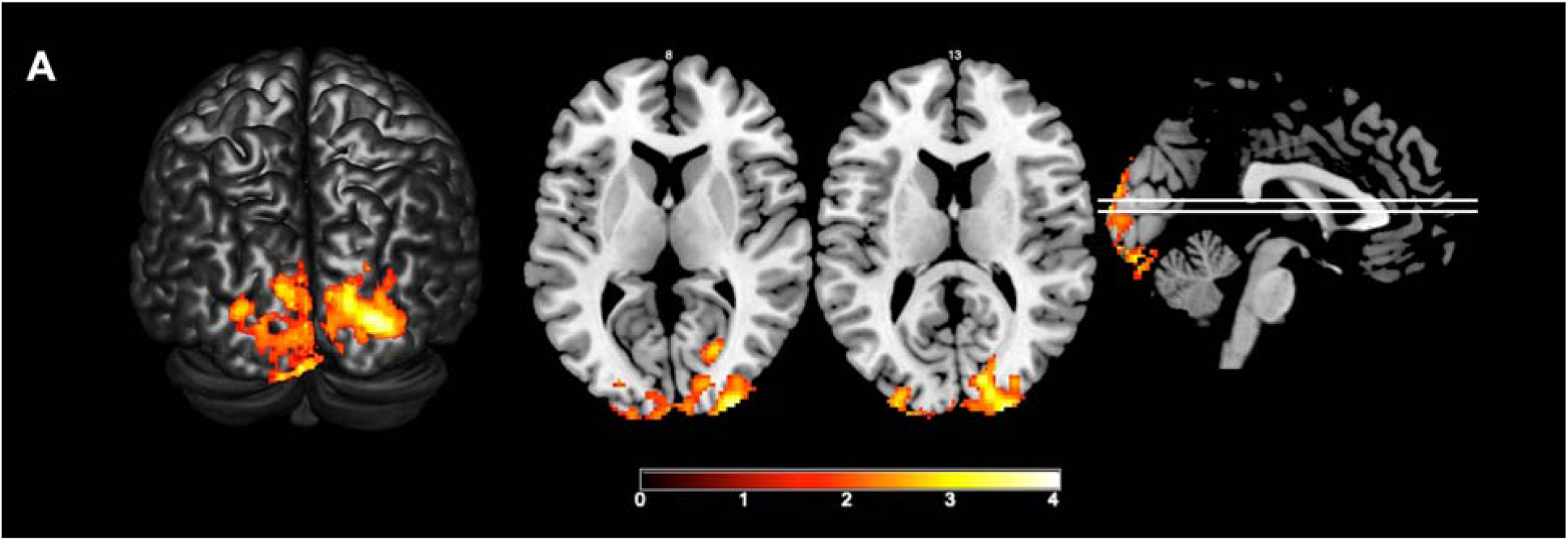
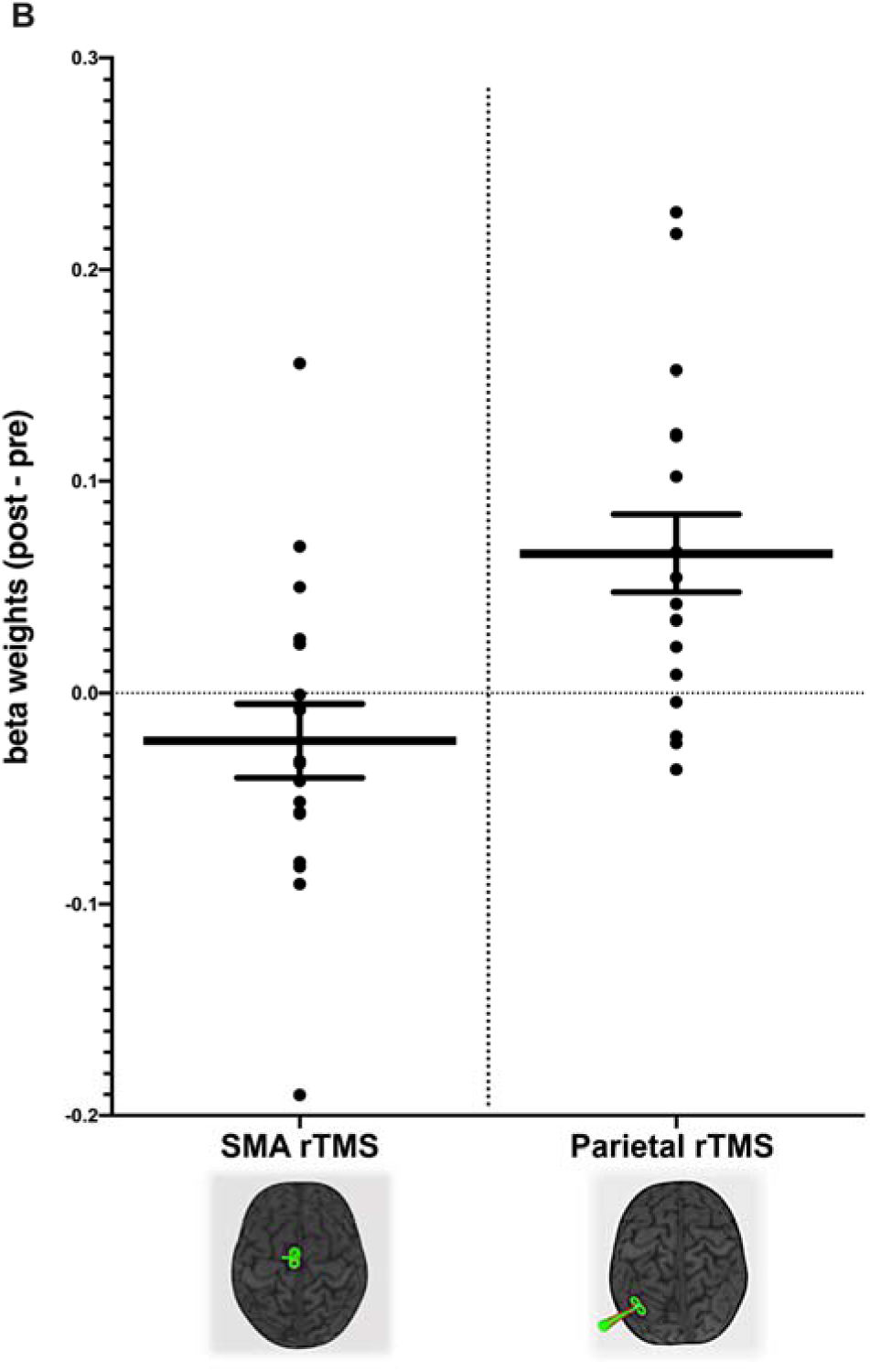
Resting-state connectivity change from pre-SMA stimulation site seed. **A)** Evidence of decreased FC between pre-SMA seed region and occipital cortex following pre-SMA stimulation, and increased FC following parietal stimulation. **B)** Beta weights obtained from occipital regions showing significant change in FC with the pre-SMA seed region following stimulation (between-subject comparison; post-pre between conditions). Left side of plot depicts changes to beta weight strength following pre-SMA stimulation, with evidence of net reduction in functional connectivity. Right side of plot depicts changes to beta weight strength following parietal stimulation, with evidence of net increase in functional connectivity. Error bars reflect mean and standard error.

Overall, we provide moderate evidence of increased connectivity between left putamen and left frontal cortex following pre-SMA stimulation. We also provide subtle evidence of differential effects on FC between the stimulation site and occipital cortex across stimulation conditions, suggesting a site-specific effect of stimulation on FC.

### 3.3 Correlation of changes in memory performance and resting-state fMRI connectivity

Previous studies have reported associations between learning and changes in functional connectivity (Sampaio-Baptista et al., 2015; Wang et al., 2014). We explored whether individual changes to associative memory performance correlated with changes to resting-state network connectivity following parietal stimulation. A Pearson’s correlation was conducted between Δ memory and Δ beta weights between stimulation site and corresponding hippocampus target for individuals who received parietal stimulation during the first week of the experiment. We could not detect a significant relationship between change in associative memory performance and change in parietal-hippocampal network connectivity, r = .25, p = .33, n = 17 (see supplementary figure S5).

## 4. Discussion

Multi-day rTMS has shown promise as a method for altering connectivity in brain networks related to memory. The aims of this study were to: 1) investigate the reproducibility of long-lasting enhancement of associative memory and FC following multi-day rTMS to the parietal cortex; and 2) assess the site specificity of any changes in memory or FC following multi-day rTMS by stimulating different sites. We applied multi-day rTMS to a region of the parietal cortex functionally integrated within a cortico-hippocampal network and to a region of the pre-SMA functionally connected with the putamen. While this study was not a direct replication of Wang et al. (2014) (we used 4 instead of 5 days of stimulation), we did not observe evidence for improvements to associative memory performance, nor increases in cortico-hippocampal FC following parietal stimulation. We did, however, observe some evidence for site-specific modulations of functional connectivity. We found divergent changes in connectivity within seeded networks following stimulation of the different cortical sites, though effects were subtle and will require independent replication.

### 4.1 Multi-day parietal rTMS, associative memory, and hippocampal connectivity

Contrary to our hypotheses, we could not find any evidence for either long-lasting enhancement of associative memory or increased parieto-hippocampal connectivity following multi-day rTMS to the parietal cortex (see figures 4-6). Instead, we found moderate evidence that multi-day rTMS did not alter associative memory performance using Bayesian statistics. Furthermore, we found no evidence of increased parieto-hippocampal connectivity in the targeted network, and subtle evidence that parietal stimulation decreased connectivity between the medial temporal lobe and stimulation sites. This finding is in contrast to previous studies following similar multi-day protocols (Wang et al., 2014; Wang & Voss, 2015). The initial study conducted by Wang et al. (2014) with N = 16, reported significant improvements in face-cued word recall accuracy following five days of stimulation to subject-specific regions of left parietal cortex. These behavioural improvements were accompanied by increased functional connectivity between the targeted parieto-hippocampal network. Impressively, a subsequent follow-up study conducted on a smaller sub-sample of the same experiment (N = 8) revealed that these effects persisted for ∼15 days following stimulation (Wang & Voss, 2015). Both the improvements in associative memory (Hermiller et al., 2019) and the increases in parieto-hippocampal connectivity (Freedberg et al., 2019b) following multi-day rTMS have recently been replicated in independent samples, although the sample sizes in these studies were moderate.

The reasons for such substantial differences between the results of our study and previous studies are unclear, however several factors may have contributed. It is possible that different sample sizes may have contributed to discrepant study outcomes. Our study was conducted using a sample of 40 healthy individuals, whereas past demonstrations of enhanced associative memory and FC have featured moderate sample sizes (N = 15-16). The response to rTMS protocols is known to be highly variable between individuals (Hamada et al., 2013; López-Alonso et al., 2014). For example, in a sample of 56 healthy individuals, Hamada et al. (2013) demonstrated that only 25% of individuals responded in the expected direction to different forms of theta burst stimulations using motor-evoked potentials as the outcome measure, while 31% responded in the opposite/ paradoxical direction. Therefore, given this pronounced inter-individual variability, it is possible that sample size differences between studies may have impacted the consistency of study outcomes. The importance of careful sample size calculations / larger samples in managing variability has been underscored by a past reviews (Guerra, López-Alonso, Cheeran, & Suppa, 2017a, 2017b). While we do not seek to question the validity of past findings, we recommend that future studies examining similar effects of multi-day rTMS and associative memory adopt comparable sample sizes to the present study.

Differences in total stimulation dosage may have influenced study outcomes. Wang and colleagues applied five daily sessions of stimulation, while in the current study we applied four sessions of stimulation utilising the same stimulation parameters (i.e. 20 Hz, 2 second trains, 28 seconds inter-trail interval). It is therefore possible that the reduced total dose applied in the current study may have attenuated stimulation-induced effects on FC. A recent study investigated the dosage utilising a Bayesian predictive approach (Freedberg et al., 2019a), concluding that a minimum of five sessions is necessary to induce increases in cortico-hippocampal FC comparable to Wang et al. (2014). However, a subsequent study by the same group found comparable increases in hippocampal FC following 3-4 sessions of parietal stimulation (Freedberg et al., 2019b). Given inconsistencies in dose-response and the absence of a physiological basis for a minimum effective dose (i.e. four vs five sessions of rTMS), further investigation of the minimum session number to elicit reliable FC changes in this context is required.

Additionally, we also acknowledge differences between our protocol and past studies in regard to the number of associative memory assessments. In the current study, associative memory assessments were conducted before and after each week of stimulation. In comparison, Wang et al. (2014) conducted an additional assessment of associative memory at a mid-week time point (i.e. prior to the third session for both active and sham stimulation conditions). This additional exposure to the face-cued word recall task, followed by stimulation shortly after (∼1 hour), may have bolstered the consolidation of associative learning, rather than inducing a generalised enhancement of associative memory capacity *per se*. The absence of this mid-week task exposure in our study may have attenuated this effect. However, a recent study has successfully replicated these associative memory effects without a mid-week assessment in a sample of 16 healthy adults (Hermiller et al., 2019), suggesting that there may also be other factors influencing our observed outcomes. It is possible that inconsistent effects of rTMS on associative memory may more broadly reflect a variable response to the stimulation. Future studies may investigate whether certain physiological or lifestyle factors (e.g. parietal-hippocampal FC at baseline or concurrent level of physical activity (Hendrikse et al. 2017; Ridding & Ziemann, 2010) are predictive of associative memory changes following rTMS.

### 4.2 Site specificity of changes in functional connectivity following multi-day rTMS

While rTMS is often applied to different cortical regions, few studies have compared the influence of stimulation site on network connectivity. Our results suggest that the effects of multi-day rTMS on FC are site-specific and differ across targeted and non-targeted networks. When seeding either the parietal stimulation site or hippocampus target within individualised cortico-hippocampal networks, we observed evidence of decreased FC following parietal stimulation and increased FC following pre-SMA stimulation. Similarly, when seeding the stimulation site of the pre-SMA-putamen network, we again observed divergent effects, with decreased FC following pre-SMA stimulation and increased FC following parietal stimulation. Thus, we report some evidence of reduced FC within targeted networks and increased FC across distinct non-targeted networks following multi-day 20 Hz rTMS. These findings align with Eldaief et al. (2011) who also reported a decrease in FC following 20 Hz rTMS to the left parietal cortex (MNI coordinate x = −46, y = −70, z = 31). Specifically, this study reported a widespread decrease in connectivity between the parietal stimulation target and interconnected regions within the default-mode network. Further, other studies have also observed site-specific changes to FC following rTMS. For example, Castrillon et al. (2020) demonstrated differential effects of 1 Hz stimulation between a posterior sensory network and a frontal cognitive network, and evidence of widespread changes to FC across non-targeted regions. A recent systematic review of the effects of rTMS on FC also concluded that rTMS-induced effects are frequently seen outside of the targeted network and overall do not follow accepted frequency-dependent conventions (i.e. low frequency (< 1 Hz) stimulation reduces FC, high frequency (> 5 Hz) increases FC) (Beynel, Powers, & Appelbaum, 2020). Overall, our findings of reduced FC within targeted networks challenges the notion that local changes to cortical excitation/inhibition induced by multi-day 20 Hz rTMS translate to similar effects at the network level.

However, there are certain limitations of our FC analysis which must be acknowledged. For example, our study conducted a between-subject comparison of FC, which may limit the generalisability of our findings. Further, given the considerable influence of pre-processing strategy on group-level FC estimates (Parkes, Fulcher, Yücel, & Fornito, 2018), we also acknowledge differences in denoising approaches between our study and those reporting increased cortico-hippocampal FC following multi-day parietal rTMS (Freedberg et al., 2019b; Wang et al., 2014). For example, we have included white matter and CSF as nuisance regressors, while past studies have not (Freedberg et al., 2019b; Wang et al., 2014). While such methodological factors can influence signal-to-noise ratio and reliability (Shirer et al., 2015), we did not reproduce this past result whether these signals were included as nuisance regressors, or not. Additionally, our reported effects were also FWE-corrected with a cluster defining height threshold of p < .05 to approximate the approach of Wang et al. (2014). When adopting a more conservative cluster defining height threshold of p < .001, these effects are absent. Further research is needed to examine the reliability of site-specific changes to connectivity following multi-day rTMS.

### 4.3 Conclusion

In summary, we could not find evidence that multi-day 20 Hz rTMS to an individualised parietal-hippocampal network improves associative memory performance, or increases FC within the targeted network. In contrast, we found subtle reductions in FC within targeted parietal networks and increases in FC within non-targeted networks at 24 hrs following stimulation. There was evidence of both increased and decreased FC within targeted pre-SMA networks. Our findings suggest a complex interplay between multi-day rTMS and network connectivity, with changes in both target and non-target networks, often in the opposite direction to those hypothesised. Future work uncovering the factors which determine how rTMS modulates cortical networks is required to develop more reliable paradigms for driving desired changes in functional connectivity between cortical and subcortical regions.

## Acknowledgements

We are grateful to Associate Professor Joel Voss and the laboratory for human neuroscience for their seminal research that has motivated this study, and for sharing the face-cued word recall task. We also wish to thank the staff at Monash Biomedical Imaging for their assistance with MRI data acquisition.

## Data and material availability

No part of the study procedures or analyses was pre-registered prior to the research being conducted. We report how we determined our sample size, all data exclusions, all inclusion/exclusion criteria, whether inclusion/exclusion criteria were established prior to data analysis, all manipulations, and all measures in the study. The face-cued word recall task materials were shared by Associate Professor Joel Voss. Requests to access these materials should be directed to the laboratory for human neuroscience (www.lhn.northwestern.edu). The conditions of our ethical approval do not permit open sharing of participant MRI data without prior informed consent. Therefore, we are unable to publicly archive the raw MRI data used in this study. De-identified behavioural data and inputs for group level fMRI analyses (i.e. the 1^st^ level fMRI contrast images) are available at https://osf.io/y2xm8/, and all code used for fMRI analysis is available at https://github.com/jhendrikse/ex_rtms_code.

## Supplementary materials

### rTMS target localisation

A consistent target localisation procedure was applied for each participant to ensure accuracy across our sample, and relative to past studies (Wang et al., 2014). Individual subject resting state connectivity profiles were generated in MNI space and the fMRI local maxima within 15mm radius of MNI coordinate x = −47, y = −68, z =36 was chosen for parietal site. The inverse transform was applied to the chosen coordinate to visually confirm the stimulation target in native space in relation to anatomical cortical landmarks. Each subject’s T1-weighted image was then used to generate a subject-specific neuronavigation profile using the Brainsight system. This process includes a series of steps in which a transformation matrix is generated that encodes the spatial warping performed on each subject’s structural for alignment into MNI space. This feature of the Brainsight software allows any coordinate to be reported in either native or standardised MNI space. The MNI target coordinate was marked onto each subject’s structural image to ensure accurate target localisation during stimulation. Again, the coordinate was visually confirmed in relation to anatomical cortical landmarks.

### Individualised sub-cortical seeds and stimulation targets

**Figure S1.**
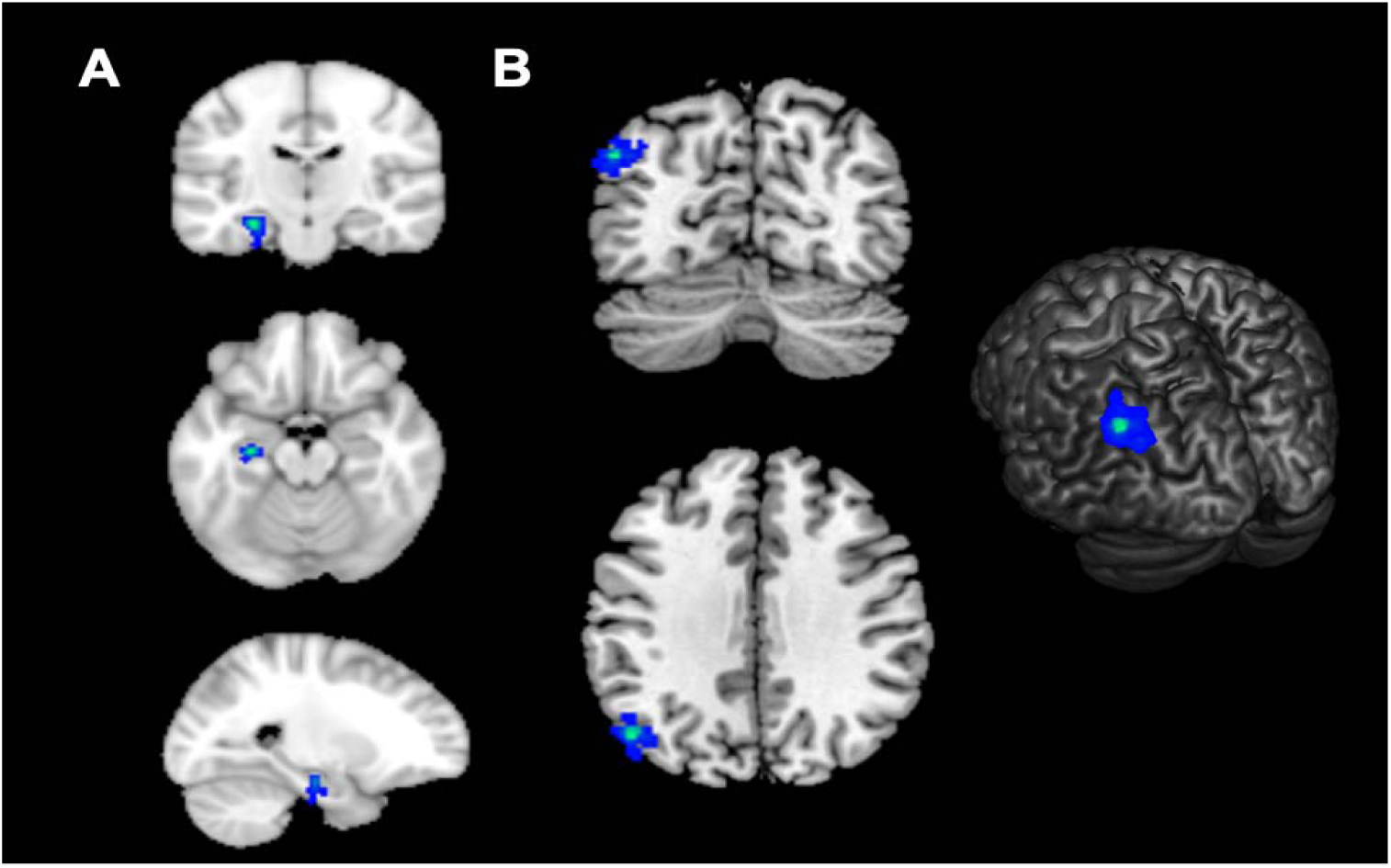
Overlay of individual hippocampus seed locations (A) and corresponding parietal stimulation targets (B), superimposed on template brain. Hippocampus seeds were localised to the middle of the hippocampus proper on each subject’s anatomical image. Parietal stimulation targets were individualised on the basis of peak functional connectivity to left hippocampus. Voxels in blue to green represent the average spatial overlap across subjects, with greater proportion of overlap represented in green.

**Figure S2.**
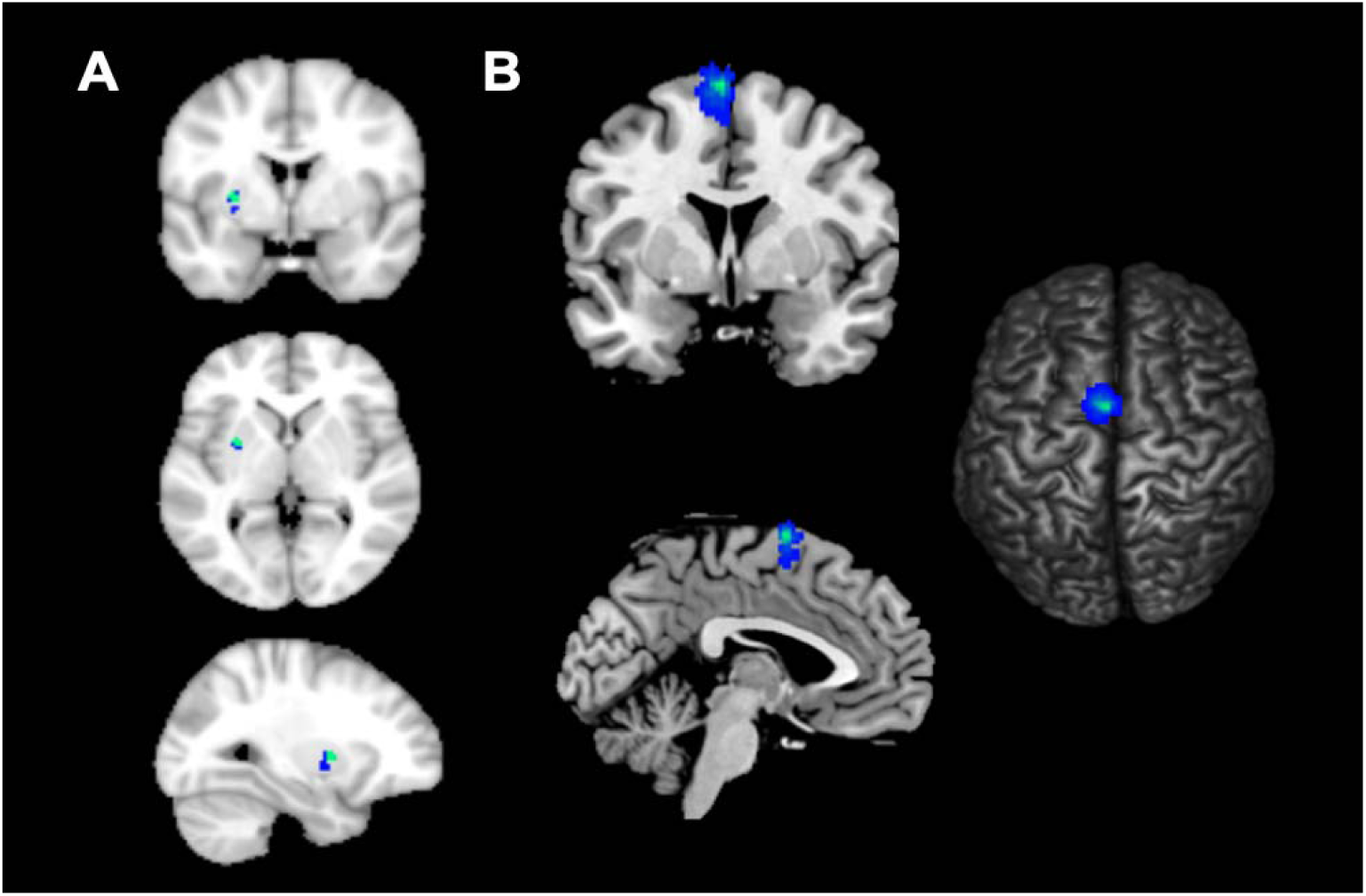
Overlay of individual putamen seed locations (A) and corresponding pre-SMA stimulation targets (B), superimposed on template brain. Putamen seed were localised to MNI coordinate x = −28, y = 1, z = 3. Small transformations in the z and y plane were performed for two subjects to mirror the anatomical locations reported by Di Martino et al. (2008). Pre-SMA stimulation targets were individualised on the basis of peak functional connectivity to left putamen.

**Figure S3.**
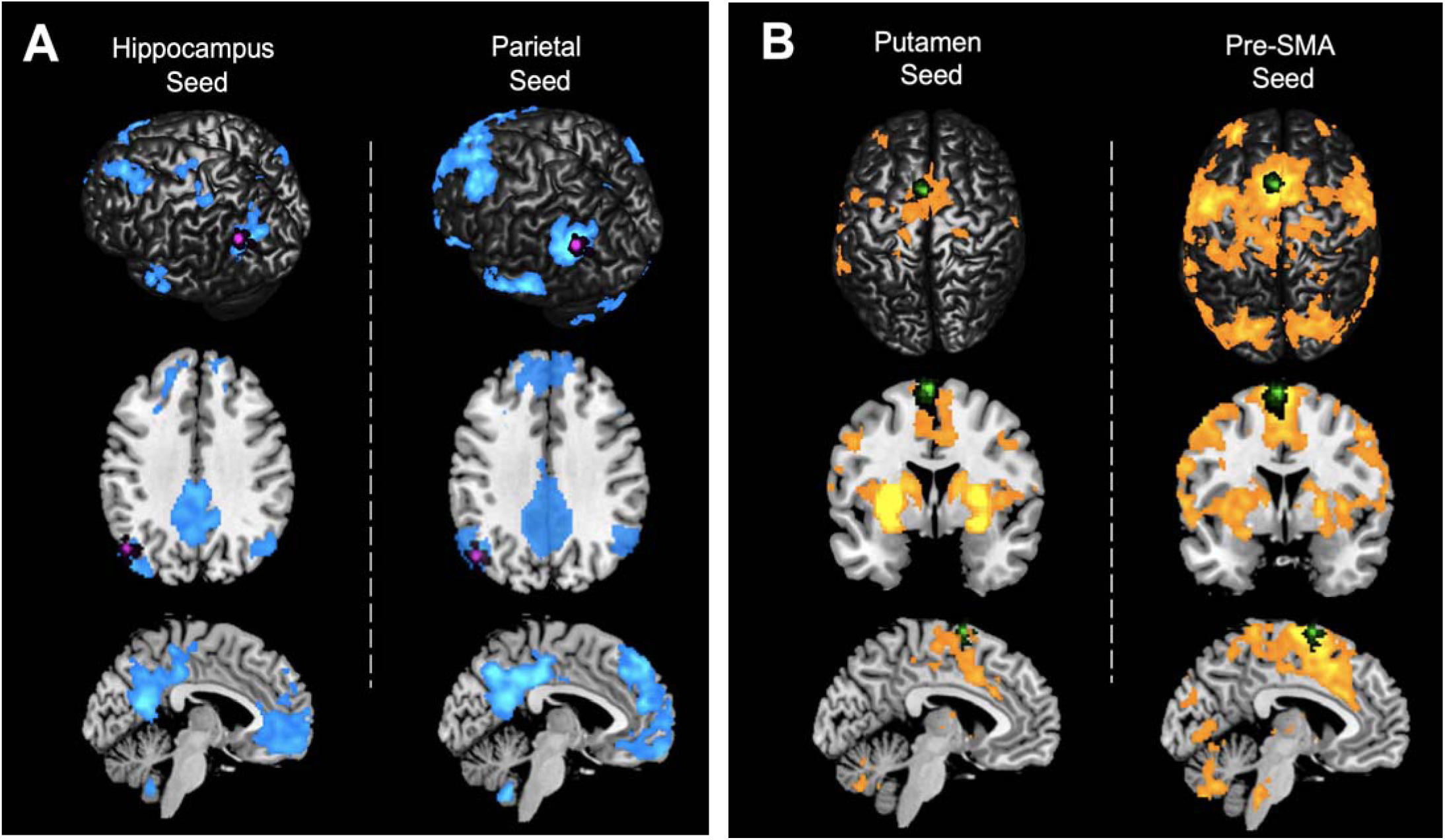
Overlay of individualised stimulation targets on averaged resting-state networks. **A)** Individual parietal stimulation targets & cortico-hippocampal resting-state network, extracted from hippocampus seed (left) and parietal stimulation site (right). **B)** Individual pre-SMA stimulation targets & cortico-putamen resting-state network, extracted from putamen seed (left) and pre-SMA stimulation site (right). Seeding either the cortical stimulation site or corresponding sub-cortical target extracted a spatially consistent network. Parietal stimulation targets are displayed in magenta and pre-SMA targets are displayed in green (voxelwise FWE p < .0001, with extent threshold k > 50 contiguous voxels for illustrative purposes).

### Analysis of associative memory changes in participations who received rTMS using the Brainsight neuronavigation system

To assess whether the use of different neuronavigation systems influenced the effects of stimulation on memory, an additional analysis was performed on the sub-sample that completed the protocol with the Brainsight neuronavigation system (N = 31). In keeping with the statistical approach presented in the manuscript, a 2 x 2 repeated-measures ANOVA (within-subject factors of Stimulation Condition and Time) was conducted on percentage correct values on the face-cued word recall task. The results revealed no significant interaction between Stimulation Condition and Time (F_1,30_ = 0.03, p = .86, *η*^2^_*p*_ = .00), main effect of Time (baseline, post-rTMS; F_1,30_ = 0.23, p = .63, *η*^2^_*p*_ = .01) or main effect of Stimulation Condition (parietal, pre-SMA; F_1,30_ = 0.92, p = .35, *η*^2^_*p*_ = .03). In summary, this analysis with neuronavigation system held constant yields the same conclusion as the main analysis i.e. multi-day rTMS had no effect on associative memory performance.

### Analysis of associative memory changes in young adult sample

Past studies examining the effects of multi-day parietal-hippocampal rTMS on associative memory have been conducted in young-adult samples (Hermiller et al., 2019; Wang et al., 2014). Our sample was predominantly comprised of young adults (35 out of 40 aged between 18-31 years), while five participants were aged between 40-55 years. To account for the possibility that young and middle-aged adults may have responded differently to multi-day rTMS, an additional analysis of associative memory was performed on the young adult sample (N = 35). A paired-sample t-test was conducted on Δ values calculated for each stimulation condition. Two outlier values (Z-score > 2.58) were identified from the pre-SMA stimulation condition, and these datapoints were removed from the analysis (as described in section 3.1). No significant differences were found between Δ parietal stimulation (M = 0.04, SD = 0.44) and pre-SMA stimulation (M = 0.01, SD = 0.44) (t_1,32_ = 0.45, p = .65, Cohen’s d = 0.08) when middle-aged participants were removed from the analysis. These results are consistent with those reported from the complete sample (N = 40).

### Comparison of associative memory performance following parietal stimulation in week one

To account for possible carryover effects of pre-SMA stimulation on the response to parietal stimulation or associative memory estimates, a paired-samples t-test was conducted on baseline and post-rTMS associative memory scores in subjects who received parietal stimulation in the first week of the study (N = 20) (i.e. subjects who had not received prior pre-SMA stimulation). There were no significant differences in associative memory present in this sub-sample with similar task performance at baseline (M = 40.00 SD = 21.76) and post-rTMS (M = 40.00, SD =24.87), t_1,19_ = 0.00, p = 1.00, Cohen’s d = 0.00, BF_10_ = 0.23.

### Analysis of network-specific connectivity between stimulation site and sub-cortical targets

#### Parietal – hippocampal connectivity

To investigate the possibility of rTMS exerting network-specific, rather than brain-wide effects, we conducted a spatially constrained analysis of functional connectivity change between *a priori* regions of interest. We expected a significant increase in beta weights representing the correlation between parietal cortex and left hippocampus following stimulation of the parietal cortex, but not following stimulation of the active control region in pre-SMA.

An independent-samples t-test of the difference in beta weight strength between parietal cortex and hippocampus following stimulation to either parietal cortex (M = −0.02, SD = 0.08) or pre-SMA active control site (M = −0.005, SD = 0.05), was not significant, t_34_ = −0.47, p = .64, Cohen’s d = 0.16. Thus, analysis of our data revealed no evidence of a change to functional connectivity between the parietal cortex stimulation site and the corresponding hippocampal target.

#### Pre-SMA – putamen connectivity

Conversely, a significant increase in beta weights between pre-SMA and left putamen were expected following pre-SMA stimulation, but not parietal stimulation.

An independent-samples t-test was also conducted to assess the effects of pre-SMA stimulation on functional connectivity between the stimulated region of the cortex and left putamen target. There was no significant difference observed in beta weight strength between pre-SMA and putamen following stimulation to either pre-SMA (M = 0.004, SD = 0.07) or parietal (M = −0.01, SD = 0.09) targets, t_34_ = 0.69, p = .49, Cohen’s d = 0.23. Thus, consistent with the cortico-hippocampal network results, we report no evidence of sustained changes to pre-SMA – putamen functional connectivity following stimulation.

**Figure S4.**
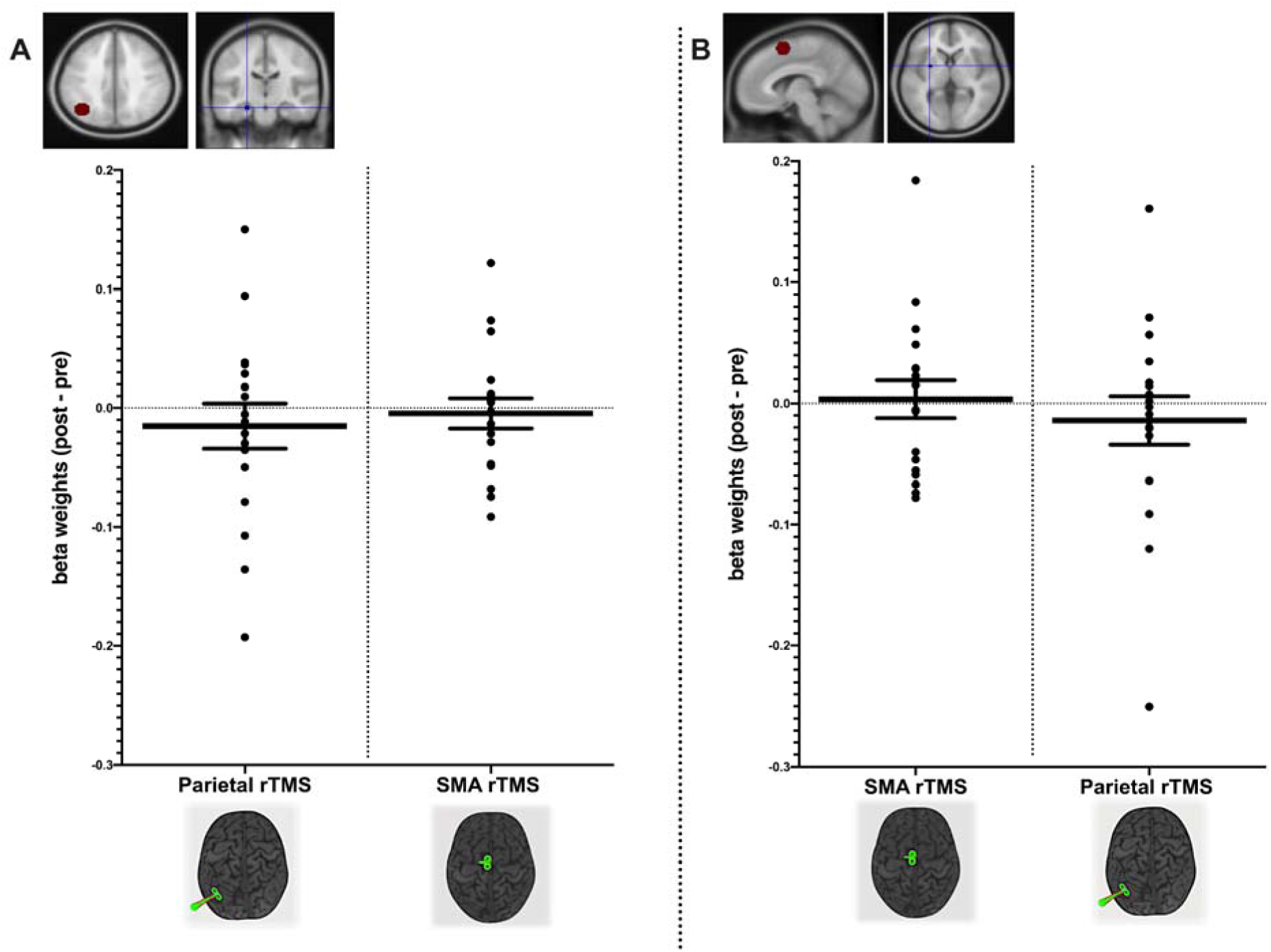
Network-specific analysis of functional connectivity change. No significant changes between stimulation site and subcortical targets. **A)** Changes to beta weight strength between left parietal cortex and left hippocampus seed following parietal stimulation (left) and pre-SMA active control stimulation (right). **B)** Changes to beta weights strength between pre-SMA and left putamen seed following pre-SMA stimulation (left) and parietal stimulation (right).

### Correlation of changes in memory performance and resting-state fMRI connectivity

We explored whether individual changes to associative memory performance correlated with changes to resting-state network connectivity following parietal stimulation. There was no significant relationship between change in associative memory performance and change in parietal-hippocampal network connectivity, r = .25, p = .33, n = 17.

**Figure S5.**
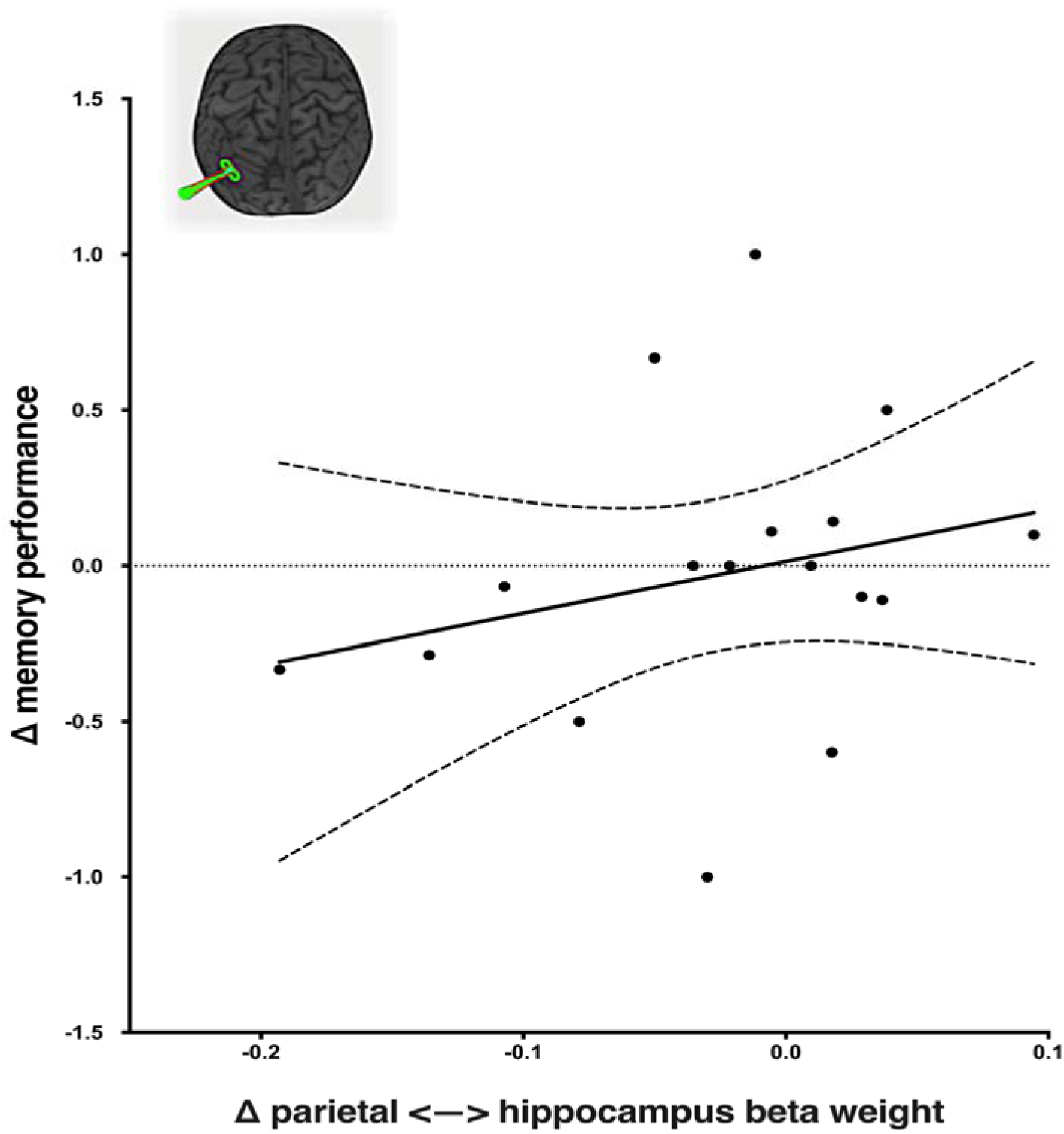
Association between changes in memory performance and resting-state fMRI connectivity. Change in associative memory following parietal stimulation (expressed as change relative to baseline performance) is shown on the y-axis and change in parietal-hippocampal connectivity (post beta weights – pre beta weights) is represented on the x-axis. Solid line shows regression line of best fit, dotted lines depict 95% confidence interval.

## References

Alexander, G. E., DeLong, M. R., & Strick, P. L. (1986). Parallel Organization of Functionally Segregated Circuits Linking Basal Ganglia and Cortex. Annual Review of Neuroscience, 9(1), 357–381. https://doi.org/10.1146/annurev.ne.09.030186.002041

Althoff, R. R., & Cohen, N. J. (1999). Eye-Movement-Based Memory Effect: A Reprocessing Effect in Face Perception. Journal of Experimental Psychology: Learning Memory and Cognition, 25(4), 997–1010. https://doi.org/10.1037/0278-7393.25.4.997

Barnes, J. M., & Underwood, B. J. (1959). “Fate” of first-list associations in transfer theory. Journal of Experimental Psychology, 58(2), 97–105. https://doi.org/10.1037/h0047507

Behzadi, Y., Restom, K., Liau, J., & Liu, T. T. (2007). A component based noise correction method (CompCor) for BOLD and perfusion based fMRI. NeuroImage, 37(1), 90–101. https://doi.org/10.1016/j.neuroimage.2007.04.042

Beynel, L., Powers, J. P., & Appelbaum, L. G. (2020). Effects of repetitive transcranial magnetic stimulation on resting-state connectivity: A systematic review. NeuroImage, 211(September 2019), 116596. https://doi.org/10.1016/j.neuroimage.2020.116596

Button, K. S., Ioannidis, J. P. A., Mokrysz, C., Nosek, B. A., Flint, J., Robinson, E. S. J., & Munafò, M. R. (2013). Power failure: Why small sample size undermines the reliability of neuroscience. Nature Reviews Neuroscience, 14(5), 365–376. https://doi.org/10.1038/nrn3475

Castrillon, G., Sollmann, N., Kurcyus, K., Razi, A., Krieg, S. M., & Riedl, V. (2020). The physiological effects of noninvasive brain stimulation fundamentally differ across the human cortex. Science Advances, 6(5). https://doi.org/10.1126/sciadv.aay2739

Cocchi, L., Sale, M. V, L Gollo, L., Bell, P. T., Nguyen, V. T., Zalesky, A., … Mattingley, J. B. (2016). A hierarchy of timescales explains distinct effects of local inhibition of primary visual cortex and frontal eye fields. ELife, 5, 1–17. https://doi.org/10.7554/eLife.15252

De Geeter, N., Crevecoeur, G., Leemans, A., & Dupré, L. (2015). Effective electric fields along realistic DTI-based neural trajectories for modelling the stimulation mechanisms of TMS. Physics in Medicine and Biology, 60(2), 453–471. https://doi.org/10.1088/0031-9155/60/2/453

Di Martino, A., Scheres, A., Margulies, D. S., Kelly, A. M. C., Uddin, L. Q., Shehzad, Z., … Milham, M. P. (2008). Functional connectivity of human striatum: A resting state fMRI study. Cerebral Cortex, 18(12), 2735–2747. https://doi.org/10.1093/cercor/bhn041

Doyon, J., Bellec, P., Amsel, R., Penhune, V., Monchi, O., Carrier, J., … Benali, H. (2009). Contributions of the basal ganglia and functionally related brain structures to motor learning. Behavioural Brain Research, 199(1), 61–75. https://doi.org/10.1016/j.bbr.2008.11.012

Draganski, B., Kherif, F., Klöppel, S., Cook, P. A., Alexander, D. C., Parker, G. J. M., … Frackowiak, R. S. J. (2008). Evidence for segregated and integrative connectivity patterns in the human basal ganglia. Journal of Neuroscience, 28(28), 7143–7152. https://doi.org/10.1523/JNEUROSCI.1486-08.2008

Eklund, A., Nichols, T. E., & Knutsson, H. (2016). Cluster failure: Why fMRI inferences for spatial extent have inflated false-positive rates. Proceedings of the National Academy of Sciences, 201602413. https://doi.org/10.1073/pnas.1602413113

Eldaief, M. C., Halko, M. A., Buckner, R. L., & Pascual-Leone, A. (2011). Transcranial magnetic stimulation modulates the brain’s intrinsic activity in a frequency-dependent manner. Proceedings of the National Academy of Sciences of the United States of America, 108(52), 21229–21234. https://doi.org/10.1073/pnas.1113103109

Esteban, O., Markiewicz, C. J., Blair, R. W., Moodie, C. A., Isik, A. I., Erramuzpe, A., … Gorgolewski, K. J. (2019). fMRIPrep: a robust preprocessing pipeline for functional MRI. Nature Methods, 16(1), 111–116. https://doi.org/10.1038/s41592-018-0235-4

Fox, M. D., Halko, M. A., Eldaief, M. C., & Pascual-Leone, A. (2012). Measuring and manipulating brain connectivity with resting state functional connectivity magnetic resonance imaging (fcMRI) and transcranial magnetic stimulation (TMS). NeuroImage, 62(4), 2232–2243. https://doi.org/10.1016/j.neuroimage.2012.03.035

Fox, N. C., Warrington, E. K., Freeborough, P. A., Hartikainen, P., Kennedy, A. M., Stevens, J. M., & Rossor, M. N. (1996). Presymptomatic hippocampal atrophy in Alzheimer’s disease. A longitudinal MRI study. Brain □: A Journal of Neurology, 119 (Pt 6(1996), 2001–2007. https://doi.org/10.1093/brain/119.6.2001

Freedberg, M., Reeves, J. A., Toader, A. C., Hermiller, M. S., Kim, E., Haubenberger, D., … Wassermann, E. M. (2019a). Optimizing Hippocampal-Cortical Network Modulation via Repetitive Transcranial Magnetic Stimulation: A Dose-Finding Study Using the Continual Reassessment Method. Neuromodulation, 2019. https://doi.org/10.1111/ner.13052

Freedberg, M., Reeves, J. A., Toader, A. C., Hermiller, M. S., Voss, J. L., & Wassermann, E. M. (2019b). Persistent enhancement of hippocampal network connectivity by parietal rTMS is reproducible. ENeuro, 6(5), 1–13. https://doi.org/10.1523/ENEURO.0129-19.2019

Galea, J. M., Albert, N. B., Ditye, T., & Miall, R. C. (2010). Disruption of the dorsolateral prefrontal cortex facilitates the consolidation of procedural skills. Journal of Cognitive Neuroscience, 22(6), 1158–1164. https://doi.org/10.1162/jocn.2009.21259

George, M. S., & Short, E. B. (2014). The Expanding Evidence Base for rTMS Treatment of Depression. Curr Opin Psychiatry, 26(1), 13–18. https://doi.org/10.1097/YCO.0b013e32835ab46d

Guerra, A., López-Alonso, V., Cheeran, B., & Suppa, A. (2017a). Solutions for managing variability in non-invasive brain stimulation studies. Neuroscience Letters, 719(October 2017), 133332. https://doi.org/10.1016/j.neulet.2017.12.060

Guerra, A., López-Alonso, V., Cheeran, B., & Suppa, A. (2017b). Variability in non-invasive brain stimulation studies: Reasons and results. Neuroscience Letters, 719(October 2017), 133330. https://doi.org/10.1016/j.neulet.2017.12.058

Hamada, M., Ugawa, Y., & Tsuji, S. (2009). High-frequency rTMS over the supplementary motor area improves bradykinesia in Parkinson’s disease: subanalysis of double-blind sham-controlled study. Journal of the Neurological Sciences, 287(1-2), 143–146. https://doi.org/10.1016/j.jns.2009.08.007

Hamada, M., Murase, N., Hasan, A., Balaratnam, M., & Rothwell, J. C. (2013). The role of interneuron networks in driving human motor cortical plasticity. Cerebral Cortex, 23(7), 1593–1605. https://doi.org/10.1093/cercor/bhs147

Heckers, S. (2001). Neuroimaging studies of the hippocampus in schizophrenia. Hippocampus, 11(5), 520–528. https://doi.org/10.1002/hipo.1068

Hendrikse, J., Kandola, A., Coxon, J., Rogasch, N., & Yücel, M. (2017). Combining aerobic exercise and repetitive transcranial magnetic stimulation to improve brain function in health and disease. Neuroscience & Biobehavioral Reviews, 83(September), 11–20. https://doi.org/10.1016/j.neubiorev.2017.09.023

Hermiller, M. S., Karp, E., Nilakantan, A. S., & Voss, J. L. (2019). Episodic memory improvements due to noninvasive stimulation targeting the cortical–hippocampal network: A replication and extension experiment. Brain and Behavior, 9(12), 1–9. https://doi.org/10.1002/brb3.1393

Héroux, M. E., Taylor, J. L., & Gandevia, S. C. (2015). The use and abuse of transcranial magnetic stimulation to modulate corticospinal excitability in humans. PLoS ONE, 10(12), 1–10. https://doi.org/10.1371/journal.pone.0144151

Jenkinson, M., Beckmann, C. F., Behrens, T. E. J., Woolrich, M. W., & Smith, S. M. (2012). FSL. NeuroImage, 62, 982–790. https://doi.org/10.1016/j.neuroimage.2011.09.015

Kahn, I., Andrews-Hanna, J. R., Vincent, J. L., Snyder, A. Z., & Buckner, R. L. (2008). Distinct cortical anatomy linked to subregions of the medial temporal lobe revealed by intrinsic functional connectivity. Journal of Neurophysiology, 100(1), 129–139. https://doi.org/10.1152/jn.00077.2008

López-Alonso, V., Cheeran, B., Río-Rodríguez, D., & Fernández-Del-Olmo, M. (2014). Inter-individual variability in response to non-invasive brain stimulation paradigms. Brain Stimulation, 7(3), 372–380. https://doi.org/10.1016/j.brs.2014.02.004

Luber, B., Kinnunen, L. H., Rakitin, B. C., Ellsasser, R., Stern, Y., & Lisanby, S. H. (2007). Facilitation of performance in a working memory task with rTMS stimulation of the precuneus: Frequency- and time-dependent effects. Brain Research, 1128(1), 120–129. https://doi.org/10.1016/j.brainres.2006.10.011

MacQueen, G. M., Campbell, S., McEwen, B. S., Macdonald, K., Amano, S., Joffe, R. T., … Trevor Young, L. (2003). Course of illness, hippocampal function, and hippocampal volume in major depression. Proceedings of the National Academy of Sciences of the United States of America, 100(3), 1387–1392. https://doi.org/10.1073/pnas.0337481100

Mansur, C. G., Myczkowki, M. L., De Barros Cabral, S., Sartorelli, M. D. C. B., Bellini, B. B., Dias, Á. M. H., … Marcolin, M. A. (2011). Placebo effect after prefrontal magnetic stimulation in the treatment of resistant obsessive-compulsive disorder: A randomized controlled trial. International Journal of Neuropsychopharmacology, 14(10), 1389–1397. https://doi.org/10.1017/S1461145711000575

Matsunaga, K., Maruyama, A., Fujiwara, T., Nakanishi, R., Tsuji, S., & Rothwell, J. C. (2005). Increased corticospinal excitability after 5 Hz rTMS over the human supplementary motor area. Journal of Physiology, 562(1), 295–306. https://doi.org/10.1113/jphysiol.2004.070755

Matthew Brett, Jean-Luc Anton, Romain Valabregue, J.-B. P. (2002). Region of interest analysis using an SPM toolbox. In Presented at the 8th International Conferance on Functional Mapping of the Human Brain. https://doi.org/10.1201/b14650-28

Miłkowski, M., Hensel, W. M., & Hohol, M. (2018). Replicability or reproducibility? On the replication crisis in computational neuroscience and sharing only relevant detail. Journal of Computational Neuroscience, 45(3), 163–172. https://doi.org/10.1007/s10827-018-0702-z

Parkes, L., Fulcher, B., Yücel, M., & Fornito, A. (2018). An evaluation of the efficacy, reliability, and sensitivity of motion correction strategies for resting-state functional MRI. NeuroImage, 171(July 2017), 415–436. https://doi.org/10.1016/j.neuroimage.2017.12.073

Ridding, M. C., & Ziemann, U. (2010). Determinants of the induction of cortical plasticity by non-invasive brain stimulation in healthy subjects. The Journal of Physiology, 588(Pt 13), 2291–2304. https://doi.org/10.1113/jphysiol.2010.190314

Rossi, S., Hallett, M., Rossini, P. M., & Pascual-Leone, A. (2009). Safety, ethical considerations, and application guidelines for the use of transcranial magnetic stimulation in clinical practice and research. Clinical Neurophysiology □: Official Journal of the International Federation of Clinical Neurophysiology, 120(12), 2008–2039. https://doi.org/10.1016/j.clinph.2009.08.016

Sale, M. V., Ridding, M. C., & Nordstrom, M. A. (2008). Cortisol Inhibits Neuroplasticity Induction in Human Motor Cortex. Journal of Neuroscience, 28(33), 8285–8293. https://doi.org/10.1523/JNEUROSCI.1963-08.2008

Sami, S., Robertson, E. M., & Miall, R. C. (2014). The time course of task-specific memory consolidation effects in resting state networks. J Neurosci, 34(11), 3982–3992. https://doi.org/10.1523/JNEUROSCI.4341-13.2014

Sampaio-Baptista, C., Filippini, N., Stagg, C. J., Near, J., Scholz, J., & Johansen-Berg, H. (2015). Changes in functional connectivity and GABA levels with long-term motor learning. NeuroImage, 106, 15–20. https://doi.org/10.1016/j.neuroimage.2014.11.032

Shafi, M. M., Westover, M. B., Fox, M. D., & Pascual-Leone, A. (2012). Exploration and modulation of brain network interactions with noninvasive brain stimulation in combination with neuroimaging. European Journal of Neuroscience, 35(6), 805–825. https://doi.org/10.1111/j.1460-9568.2012.08035.x

Shirer, W. R., Jiang, H., Price, C. M., Ng, B., & Greicius, M. D. (2015). Optimization of rs-fMRI Pre-processing for Enhanced Signal-Noise Separation, Test-Retest Reliability, and Group Discrimination. NeuroImage, 117, 67–79. https://doi.org/10.1016/j.neuroimage.2015.05.015

Song, X. W., Dong, Z. Y., Long, X. Y., Li, S. F., Zuo, X. N., Zhu, C. Z., … Zang, Y. F. (2011). REST: A Toolkit for resting-state functional magnetic resonance imaging data processing. PLoS ONE, 6(9). https://doi.org/10.1371/journal.pone.0025031

Stagg, C. J., Wylezinska, M., Matthews, P. M., Johansen-Berg, H., Jezzard, P., Rothwell, J. C., & Bestmann, S. (2009). Neurochemical effects of theta burst stimulation as assessed by magnetic resonance spectroscopy. Journal of Neurophysiology, 101(6), 2872–2877. https://doi.org/10.1152/jn.91060.2008

Thielscher, A., Opitz, A., & Windhoff, M. (2011). Impact of the gyral geometry on the electric field induced by transcranial magnetic stimulation. NeuroImage, 54(1), 234–243. https://doi.org/10.1016/j.neuroimage.2010.07.061

Wagner, A. D., Shannon, B. J., Kahn, I., & Buckner, R. L. (2005). Parietal lobe contributions to episodic memory retrieval. Trends in Cognitive Sciences, 9(9), 445–453. https://doi.org/10.1016/j.tics.2005.07.001

Wang, J X, Rogers, L. M., Gross, E. Z., Ryals, A. J., Dokucu, M. E., Brandstatt, K. L., … Voss, J. L. (2014). Targeted enhancement of cortical hippocampal brain networks and associative memory. Science, 345(6200), 1054–1057. https://doi.org/10.1126/science.1252900

Wang, J. X, & Voss, J. L. (2015). Long-lasting enhancements of memory and hippocampal-cortical functional connectivity following multiple-day targeted noninvasive stimulation. Hippocampus, 25(8), 877–883.

Warren, K. N., Hermiller, M. S., Nilakantan, A. S., & Voss, J. L. (2019). Stimulating the Hippocampal posteriormedial network enhances task-dependent connectivity and memory. ELife, 8, 1–21. https://doi.org/10.7554/eLife.49458

Willner, P., Scheel-Krüger, J., & Belzung, C. (2013). The neurobiology of depression and antidepressant action. Neuroscience and Biobehavioral Reviews, 37(10), 2331–2371. https://doi.org/10.1016/j.neubiorev.2012.12.007

